# Allosteric modulation of protein kinase A in individuals affected by NLPD-PKA, a neurodegenerative disease in which the RIβ-L50R variant is expressed

**DOI:** 10.1101/2024.06.30.601371

**Authors:** Tal Benjamin-Zukerman, Valeria Pane, Rania Safadi-Safa, Meir Solomon, Varda Lev-Ram, Mohammad Aboraya, Anwar Dakwar, Daniela Bertinetti, Andrew Hoy, Merel O Mol, John van Swieten, Rodrigo Maillard, Friedrich W Herberg, Ronit Ilouz

## Abstract

Protein kinase A (PKA) is a crucial signaling enzyme in neurons, with its dysregulation being implicated in neurodegenerative diseases. Assembly of the PKA holoenzyme, comprising a dimer of heterodimers of regulatory (R) and catalytic (C) subunits, ensures allosteric regulation and functional specificity. Recently, we defined the RIβ-L50R variant as a causative mutation that triggers protein aggregation in a rare neurodegenerative disease. However, the mechanism underlying uncontrolled PKA allosteric regulation and its connection to the functional outcomes leading to clinical symptoms remain elusive. In this study, we established an *in vitro* model using patient-derived cells for a personalized approach and employed direct measurements of purified proteins to investigate disease mechanisms in a controlled environment. Structural analysis and circular dichroism spectroscopy revealed that cellular proteins aggregation resulted from misfolded RIβ-subunits, preventing holoenzyme assembly and anchoring through A Kinase Anchoring Proteins (AKAPs). While maintaining high affinity to the C subunit, the resulting RIβ-L50R:C heterodimer exhibits reduced cooperativity, requiring lower cAMP concentrations for dissociation. Consequently, there was an increased translocation of C-subunit into the nucleus, impacting gene expression. We successfully controlled C subunit translocation by introducing a mutation that decreased RIβ:C dissociation in response to elevated cAMP levels. This research thus sets the stage for developing therapeutic strategies that modulate PKA assembly and allostery, thus exerting control over the unique molecular signatures identified in the disease-associated transcriptome profile.

## Introduction

Protein Kinase A (PKA) is an important signaling enzyme that plays a fundamental role in diverse neuronal functions including synaptic plasticity, neuronal development, dopamine synthesis, and long-term memory (1–5). PKA dysfunction contributes to the cognitive decline observed in many neurodegenerative diseases such as Alzheimer’s disease (AD), Parkinson’s disease (PD) and amyotrophic lateral sclerosis (ALS) (6–8). In AD, PKA is implicated in neurofibrillary pathology, where increased tau phosphorylation by PKA leads to tau aggregation and toxicity (9, 10). Modulation of Leucine-rich repeat kinase 2 (LRRK2) activity through PKA phosphorylation is linked to the progression of PD (11, 12), while reduced PKA activity is seen in the spinal cords of patients with ALS due to loss of PKA-rich motor neurons (13). Moreover, abnormal PKA pathway regulation is also observed in Huntington disease progression (14).

PKA function is regulated by the assembly of isoform-specific holoenzymes localized to specific cellular sites (15). In the inactive state, PKA exists as a holoenzyme that comprises a regulatory (R) subunit dimer and two catalytic (C) subunits which together are assembled into a quaternary structure (R_2_:C_2_) (16). Three catalytic isoforms of (Cα, Cβ, Cγ) are known, as are four regulatory subunits (RIα, RIβ, RIIα, and RIIβ). The crystal structures of RIα_2_:C_2_, RIβ_2_:C_2_, and RIIβ_2_:C_2_ revealed how isoform-specific assembly can create distinct quaternary structures that allow functional non-redundancy of the regulatory isoforms (16–18). The various PKA holoenzymes differ in terms of their allosteric mechanism and sensitivity to cAMP, as well as their putative associations with diseases resulting from the appearance of the R-subunit variants. While the incorporation of different R-subunits is linked to various maladies, only mutations in the PRKAR1B gene, encoding for the RIβ subunit, are linked with neurodevelopmental or neurodegenerative diseases (19, 20). Factional specificity of RIβ is provided by the precise localization of this subunit within various brain regions (17). PKA holoenzymes are typically targeted at specific microdomains within the cell via interaction of the R subunit with A Kinase Anchoring Proteins (AKAPs) (21).

All R subunits share similar multi-domain organization. Each R subunit consists of a Dimerization and Docking (D/D) domain in the N-terminal region, followed by an inhibitor sequence, and two tandem cyclic nucleotide binding domains (referred to as CNB-A and CNB-B) that binds cAMP and activates the enzyme. The D/D domain forms an isologous dimeric structure crucial for assembling the PKA holoenzyme into a dimer comprising two R:C heterodimers. Additionally, the D/D domain functions as a docking site for AKAPs. Given the central role of the D/D domain in maintaining structural integrity of the PKA holoenzyme, this region is evolutionary conserved (22, 23). In contrast to the abundance of known variants affecting the CNB domains, only a few variants affecting this region have been observed. Strikingly, such variants are associated with pathogenic effects (20, 24).

Recently, we defined the RIβ-L50R variant as a causative agent driving an age-dependent behavioral and neurodegenerative disease phenotype in human and mouse models. We proposed the name Neuronal Loss and Parkinsonism Driven by a PKA mutation (NLPD-PKA) for this condition, given how it encapsulates the key pathological features observed in individuals with this disease. Mechanistically, this mutation disrupts RIβ dimerization, leading to RIβ aggregation of its monomers in both human and mouse brains (25).

In the present study, we investigated several pathogenic RIβ variants presenting mutations within the D/D domain, defining RIβ-L50R as an exclusive variant able to disrupt RIβ dimerization and the assembly of PKA holoenzyme as demonstrated with cells derived from patient expressing the heterozygous L50R mutant. Furthermore, we observed abnormal cooperativity in PKA holoenzyme activation using recombinant proteins. Comprehensive analysis relying on structural, biophysical, biochemical, and cellular approaches revealed how the L50R-encoding variation destabilizes the D/D domain, resulting in protein misfolding and increased oligomerization. Specifically, we found that RIβ-L50R forms a heterodimer (RIβ:C), rather than a dimer of heterodimers (RIβ_2_:C_2_), with increased sensitivity to cAMP. Such heterodimer dynamics dysregulation was reflected in altered RNA expression patterns, thus providing a signature of neurodegenerative disease driven by the RIβ-L50R variant.

## Results

### Structural analysis of RIβ subunit variants

The RIβ subunit comprises a dimerization and docking (D/D) domain, followed by an inhibitor sequence and two cAMP binding domains. Fig. 1A highlights all identified human RIβ variants, found at the UniProt database. Variants that have been identified as likely being pathogenic or confirmed as pathogenic are denoted. In our analysis, we identified a pattern to the RIβ variants, with changes being primarily clustered within the cAMP-binding domains. In contrast, the D/D domain contained fewer such differences. The limited distribution of modified residues suggests that conservation of the RIβ sequence is fundamental for maintaining the structural integrity and functional stability of the holoenzyme. To understand the pathological consequences of genetic diversity within the *PRKAR1B*-encoding RIβ D/D domain and its influence on PKA regulatory mechanisms, we focused our analysis on four specific variants, namely, the I40V, L50R, A67V, R68Q (positions indicated with red circles in Fig. 1A).

**Figure 1.**
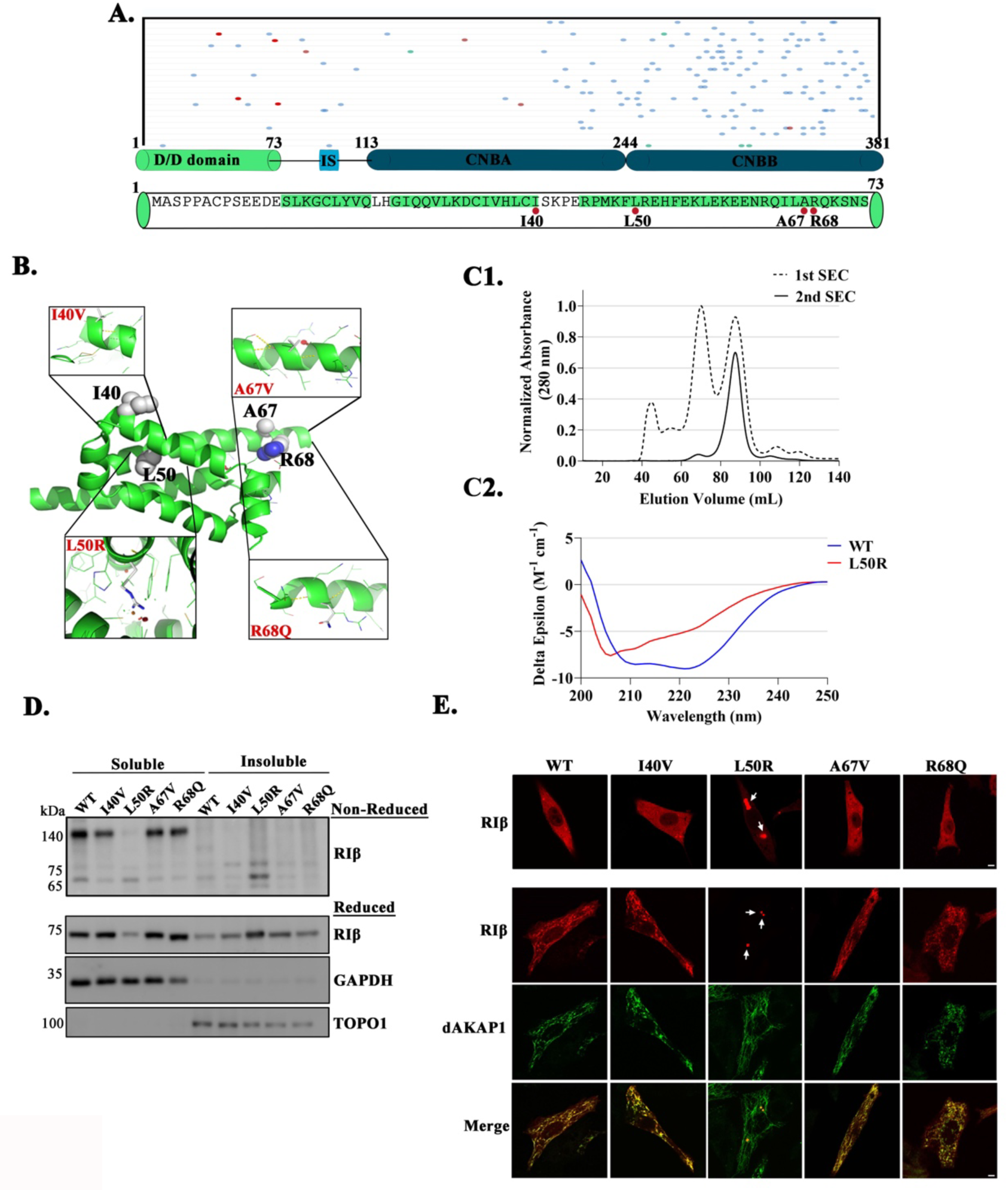
PRKAR1B variants. A. PRKAR1B domain structure encompasses a dimerization and docking (D/D) domain, succeeded by an inhibitor sequence and a pair of cAMP binding domains. Displayed below are all the identified variants of human PRKAR1B, sourced from the UniProt database. Variants that have been identified as likely pathogenic or confirmed pathogenic are marked with circles. Red circles pinpoint the variants that are the focus of this manuscript. The sequence for the D/D domain is presented, with the residues being investigated in this work accentuated. Helices A0, A1, and A3 from each protomer contribute to dimerization. **B.** In each R-subunit, the D/D domain establishes an isologous dimer. This arrangement features identical binding sites on both subunits that interface complementarily when rotated by 180° in relation to one another. Disease-related residues are represented as spheres. Detailed views of individual residues under investigation are shown in boxes. **C1.** After its purification, the 1^st^ size-exclusion chromatography (SEC) elution profile for the D/D domain displays multiple peaks likely corresponding to multimeric species. The dimeric D/D is expected to elute at ∼90 mL. Collection of ∼90 mL peak and re-injection to SEC does not show repopulate of multimeric species, indicating that the dimeric D/D domain is stable. **C2.** Circular dichroism (CD) spectrum of the wild-type D/D domain displays two minima at 208nm and 222nm, which is characteristic of α-helical proteins. The CD spectrum of L50R loses the typical features of α-helices, indicating that this mutation disrupts the D/D domain fold. Deconvolution of the CD data indicates that the wild-type and L50R proteins have 87% and 43% helicity, respectively). **D.** PC12 cells were co-transfected to express mKO2-tagged RIβ-WT or RIβ-variants and mCerulean-dAKAP1. Cell lysates divided into soluble and insoluble fractions which were separated by SDS-PAGE under both reduced (upper gel) or non-reduced conditions (lower gel). GAPDH served as a loading control for the soluble fraction, while TOPO1 was the loading control for the insoluble fraction **E.** Confocal microscopy images depict transiently transfected PC12 cells expressing mKO2-tagged RIβ-WT or mutants (upper panel) and the same cells co-transfected using mKO2-tagged RIβ-WT or mutants and dAKAP1(lower panel). Representative Images were captured at 63X magnification, with arrows highlighting the aggregates. The scale bar represents 5 µm.

We conducted structural analysis and *in-silico* modeling to investigate these four amino acid substitutions. Employing the PyMOL Mutagenesis Wizard, we implemented these replacements on the solved D/D domain structure (structure PDB ID: 4F9K). Dimerization of RIβ occurs when two monomers interlock to form a symmetrical structure consisting of three α-helices (structure PDB ID:4F9K). Within each monomer, hydrophobic and polar residues cooperate to maintain dimerization. The hydrophobic groove formed by these two protomers serves as a docking site for AKAPs. We hypothesized that RIβ variants I40V, R68Q and A67V, in which the modified residues are exposed to the surrounding solution might not be affected in terms of dimer formation nor AKAP binding (Fig. 1B). However, during the *in-silico* modeling process of RIβ variants, where we made efforts to minimize clashes, it became evident that in the L50R replacement could not avoid clashes with the surrounding residues, indicating that dimer formation could be disrupted by this mutant (Fig. 1B). This finding emphasizes the potential significance of the evolutionary conserved L50 in preserving dimerization and hence, the integrity of holoenzyme assembly, subsequently allowing for allosteric regulation. To validate the *in-silico* modeling, we used CD spectroscopy to study the secondary structure of the native RIβ and L50R variant D/D domains. Purified dimeric native RIβ D/D domain (Fig. 1C) has a typical CD spectrum for an α-helical protein, with two minima at 208 nm and 222 nm. However, L50R displayed loss of absorption at both 208nm and 222 nm, indicating a dramatic loss of α-helices. In fact, a quantitative analysis of the CD spectra revealed that RIβ wild type (RIβ -WT) has 87% of α-helices (in agreement with the high-resolution structure PDB ID: 4F9K) whereas the mutant has 43%. Thus, the loss of α-helical content for L50R provides evidence by which the full-length protein is unable to form dimers in solution like RIβ-WT. We found that the mutation disrupts the secondary structure of the three α-helices that forms the D/D domain fold.

### Impact of PRKAR1B variants on R-subunit homodimerization in the cell

To validate the structural results, we performed biochemical and cellular analyses on mutant RIβ proteins. Accordingly, we introduced DNA encoding the I40V, R68Q, A67V or L50R variants into a mKO2 fluorescently tag-encoding vector and transiently over-expressed the RIβ-WT or mutant RIβ in dissociated PC12 cells. After resolving the soluble and insoluble protein fractions from transformed cell lysates and separating their contents by SDS-PAGE under non-reducing conditions, RIβ-WT and the I40V, R68Q, A67V mutants were detected as dimers in the soluble fractions. The L50R variant, however, was found in the insoluble fraction as monomers (Fig. 1D, gel denoted as non-reduced). When the same proteins were separated by SDS-PAGE under reducing conditions, it was noted that only the L50R replacement caused RIβ to aggregate as it was found in the insoluble fraction. The other RIβ mutant proteins were mostly soluble (Fig. 1D, gel denoted as reduced). In these experiments, GAPDH and TOPO1 were used as markers to ensure efficiency of separation of the soluble and insoluble fractions, respectively.

### Impact of RIβ variants on RIβ-dAKAP1 interaction and sub-cellular localization

To further investigate the RIβ variants and their subcellular localization when expressed alone or in the presence of dAKAP1, we co-transfected PC12 cells to individually express mKO2-tagged RIβ-WT or the I40V, R68Q, A67V, or L50R RIβ variants (Fig. 1E, upper images) or together with mCerulean-tagged dAKAP1 (Fig. 1E, lower images; red for RIβ variants, green for dAKAP1). Consistent with the biochemical results presented in Fig. 1D, the RIβ protein variants in the soluble fraction exhibited a diffused cytoplasmic pattern in the immunofluorescence (IF) images. Conversely, the RIβ-L50R variant detected in the insoluble fraction was observed in aggregates in the IF images (Fig. 1E, upper images). Due to its impaired dimerization, we hypothesized that the L50R variant disrupted the co-localization of the RIβ proteins with dAKAP1 protein in the mitochondria, given how the dAKAP1-binding site is found in the dimerization interface. On the other hand, although the modified I40V, R68Q and A67V residues are located at the hydrophobic groove, our *in-silico* modeling did not predict any interreference with AKAP binding, despite their proximity to the crucial binding site. As anticipated, the co-expression of the native protein or the I40V, R68Q, A67V RIβ protein variants with dAKAP1 led to the recruitment of RIβ to the mitochondria (Fig. 1E). However, in accordance with our prediction, RIβ-L50R did not co-localize with dAKAP1 at the mitochondria. These results emphasize the importance of the L50 residue in influencing the interaction between RIβ and dAKAP1, revealing the molecular intricacies of this association.

### Patient-derived cells as an *in vitro* model for the neurodegenerative disease NLPD-PKA

To investigate the molecular and cellular mechanisms of the disease associated with the RIβ-L50R variant, i.e. NLPD-PKA, we performed analyses using primary fibroblasts derived from skin biopsies of both a healthy individual and a patient heterozygous for the mutation. These cells capture the chronological and biological aging aspects of the patients. The cells were obtained from a female patient aged 54 years for whom we had previously presented MRI scans and clinical symptoms associated with this individual, along with an age- and sex-matched healthy individual (25). We initially performed biochemical analyses on cell lysates extracted from these primary cells. The lysates were loaded and separated by SDS-PAGE under reduced conditions to determine the ratio between monomeric and dimeric forms of RIβ, as well as to evaluate the total expression levels of RIβ and C subunits. In the patient expressing the RIβ-L50R variant, expression levels of RIβ proteins in the lysates were lower than in those from the healthy control. Furthermore, there was reduced presence of RIβ proteins in their dimeric form, as compared to the healthy individual (Fig. 2A-C). The distribution between monomers and dimers was quantified based on Coomassie-stained SDS-PAGE band intensities (Fig. 2B). Importantly, the *PRKAR1B* mutation leading to the appearance of the RIβ-L50R variant did not impact C subunit expression levels of (Fig. 2D). GAPDH was used to ensure equal protein loading on the gels (Fig. 2A). Considering that the dimerization domain is folded into an isologous dimer stabilized by inter-disulfide bonds between the two protomers, we also conducted SDS-PAGE under non-reducing conditions to preserve any dimers. In the patient-derived sample, there was a significant reduction in the level of RIβ in their dimeric form, as compared to the sample from the healthy individual (Fig. 2E-F).

**Figure 2.**
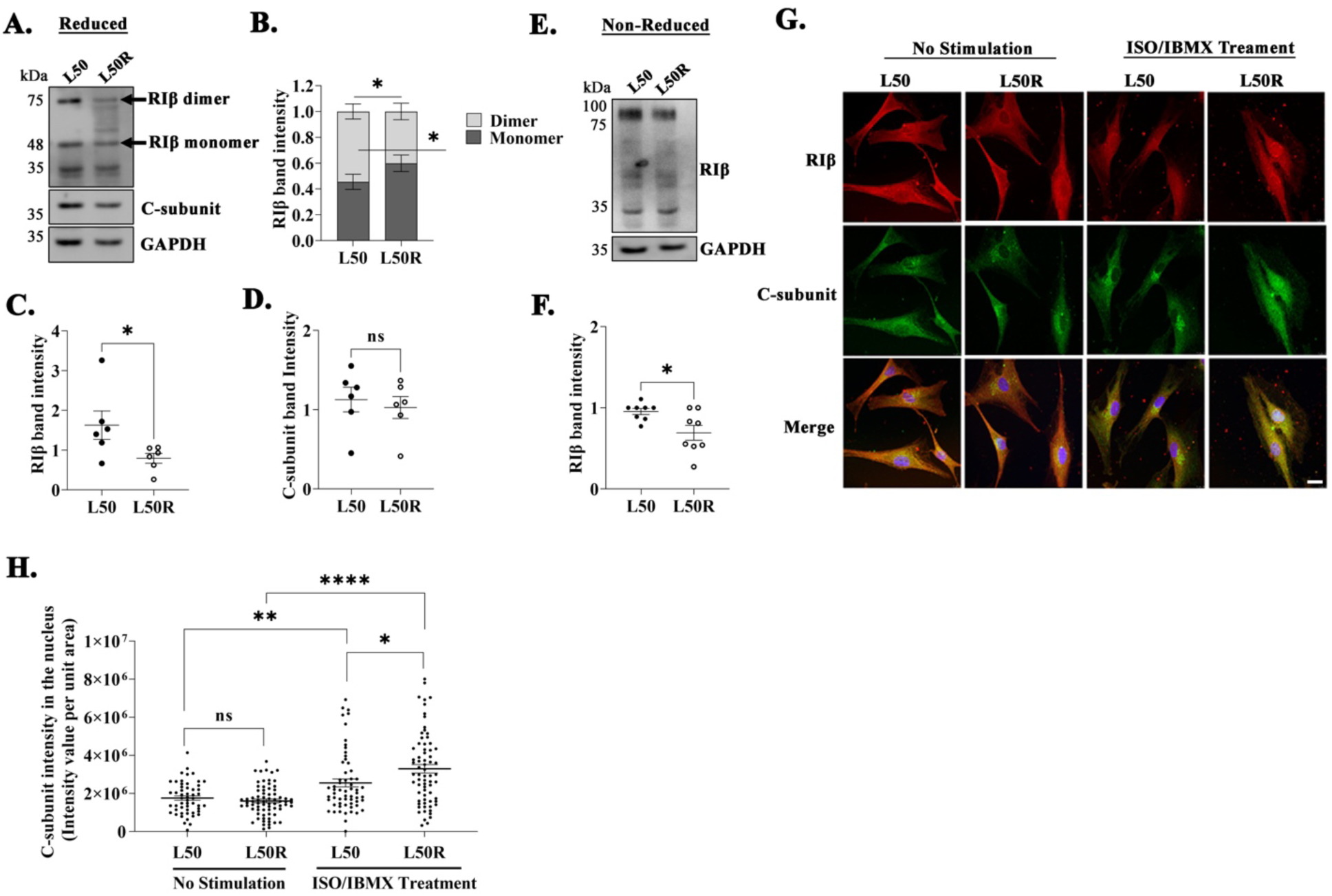
Impact of the L50R heterozygous mutation on RIβ dimerization and C-subunit nuclear translocation in patient-derived cells. **A.** Comparison of RIβ protein expression in cell lysates from a healthy individual (L50) and a patient with the L50R heterozygous mutation. Lysates were subjected to SDS-PAGE under reducing conditions. Arrows indicate the migration of RIβ proteins in both monomer and dimer forms. The C-subunit expression was determined using a specific antibody. GAPDH was used as a loading control. **B.** Quantification of RIβ band intensities from six separate experiments to determine the ratio of RIβ monomers to dimers. Mann-Whitney test was performed *P<0.05. **C.** Overall expression levels of RIβ proteins were measured by quantifying band intensities from 6 independent experiments. Mann-Whitney test was performed *P<0.05. **D.** Quantification of C-subunit expression based on band intensity from 6 separate experiments. Mann-Whitney test was performed. ns;non-significant **E.** SDS-PAGE under non-reducing conditions was conducted to compare total expression levels of the dimeric form of RIβ between L50 and L50R patients. **F.** Total expression of RIβ protein expression from the gel run in E was quantified based on band intensity from 6 independent experiments. Mann-Whitney test was performed *P<0.05. **G.** Imaging of patient-derived cells was done both pre- and post-treatment. The treatment consisted of 1 µM Isoproterenol (ISO) and 200 µM IBMX. Control treatment includes DMSO. Representative images were captured at 63X magnification 30 minutes post-treatment. Scale bar: 5 µm. **H.** Quantification of C-subunit intensity in the nucleus (total intensity per unit area) before and after cell treatment with isoproterenol and IBMX treatment for 30 min. Three independent experiments were conducted. Each dot present one cell. Error bars represents ±SEM, unpaired t test. *P<0.05, **P<0.01, ****P<0.0001, ns;non-significant.

### The RIβ-L50R variant promotes C-subunit translocation into the nucleus

The presence of RIβ-L50R in the monomeric rather than the dimeric form, combined with the unchanged expression levels of the C-subunit, led us to consider the sub-cellular localization of these subunits in the primary fibroblasts of both a healthy (L50 in Fig. 2G- H) and a patient carrying the mutation in *PRKAR1B* (L50R in Fig. 2G-H). Without cell stimulation that affects cAMP levels, RIβ and C-subunit were similarly localized of in both individuals. Following treatment with isoproterenol (ISO) and Isobutylmethylxanthine (IBMX) to raise cAMP levels, an increased concentration of C-subunit in the nucleus was noted in the healthy individual. However, the elevation in the expression of C-subunit in the patient expressing the mutant protein was significantly higher (Fig. 2G-H). These results suggest that in the presence of the RIβ-L50R variant, the L50R:C heterodimer dissociates more readily when cAMP levels in cells increase.

### Controlling rapid Cα subunit translocation into the nucleus by introducing the R211K replacement into the RIβ-L50R:Cα heterodimer

To further explore Cα translocation into the nucleus in response to cAMP-induced RIβ:Cα complex dissociation, fluorescently labeled Cα and RIβ-WT or mutant RIβ were co-expressed in PC12 cells. Confocal microscopy was utilized to observe cellular dynamics before and after treatment with forskolin (FSK) and IBMX for 30 and 60 min, treatments which strongly elevate cAMP levels and causes dissociation of the RIβ:C heterodimer complex. An additional mutation that encodes a R211K replacement in the cAMP-binding site A was introduced into the RIβ-L50R-expressing construct. This mutation is known to reduce R subunit sensitivity to cAMP (26). PC12 cells co-expressing RIβ-WT or RIβ-L50R or RIβ-L50R-R211K along with Cα were examined with or without cAMP level-affecting stimulation (Fig. 3). Consistent with findings in the patient-derived cells expressing the RIβ-L50R mutant, an increase in Cα translocation was observed upon such treatment of cells over-expressing the RIβ-L50R mutant after 30 min, as compared to the 60 min of treatment of cells over-expressing RIβ-WT required to realize the same degree of Cα translocation, indicating easier dissociation of the RIβ-L50R:C complex than of the native complex (Fig. 3A-C). The introduction of R211K into RIβ-L50R led to a mutant which failed to respond to elevated cAMP levels, and the failure of Cα to translocate into the nucleus (Fig. 3A-C). To further elucidate the sensitivity of the RIβ-L50R:C complex to cAMP and to assess the effect of the R211K replacement on Cα translocation, we performed live cell imaging. Fluorescence intensity of the nucleus was calculated as an average intensity from images taken from five regions of the plate containing the labelled cells in the same time window.

**Figure 3.**
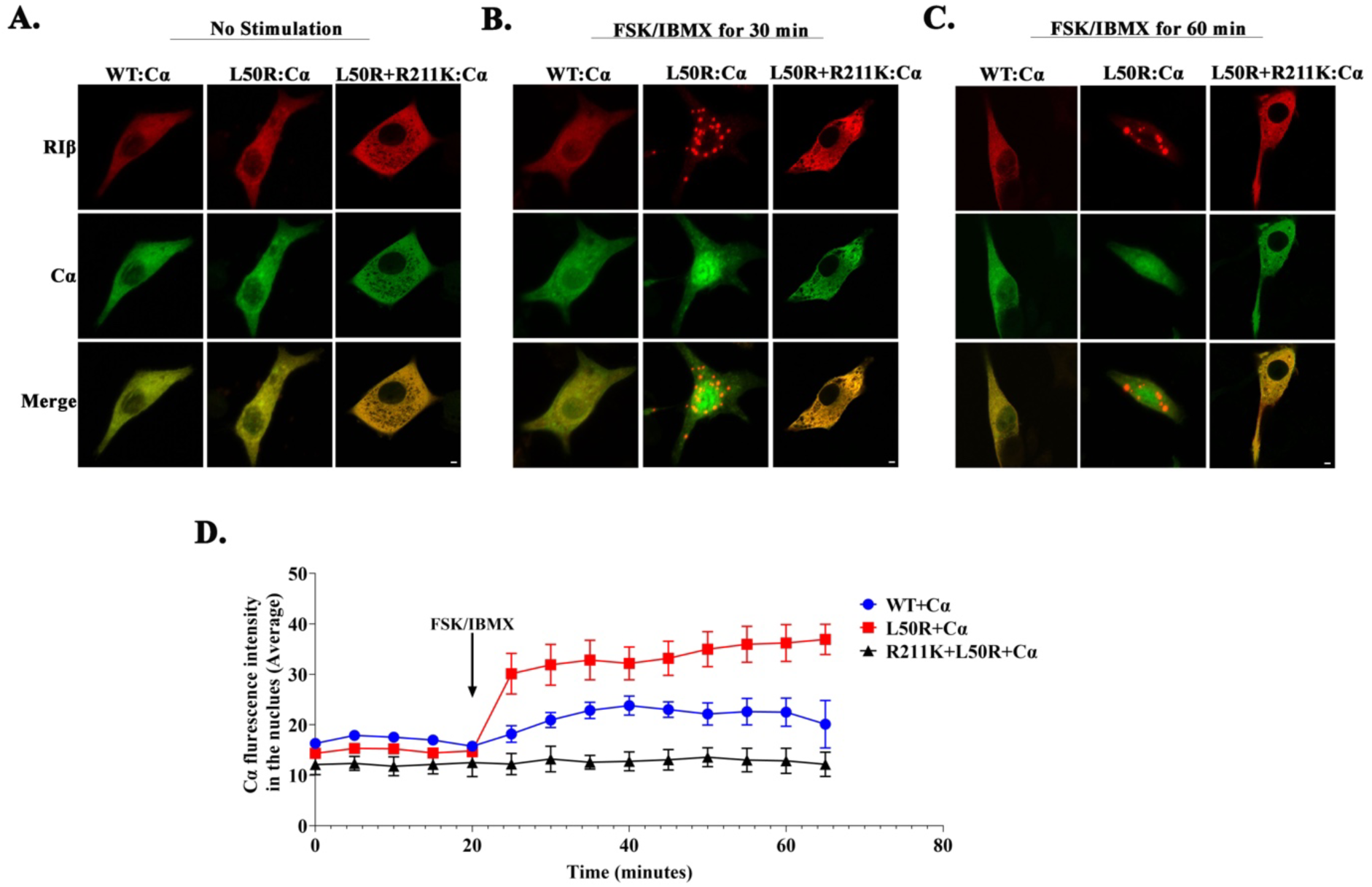
Controlling rapid Cα-subunit translocation into the nucleus by introducing the R211K mutation into the RIβ-L50R:Cα heterodimer. PC12 cells were co-transfected to express mCerulean-Cα along with mKO2-RIβ-WT or mKO2-RIβ-L50R or mKO2-RIβ-L50R+R211K. Cells were imaged before treatment (**A**), and after treatment with 20 µM Forskolin (FSK) and 200 µM IBMX for 30 min (**B**) or 60 min (**C**) post fixation. Representative images taken at 63X magnification are depicted. **D.** Live cell imaging of PC12 cells co-transfected to express mCerulean-Cα along with RIβ-WT or RIβ-L50R or RIβ-L50R+R211K. The cells imaged every 5 min for 20 min before treatment and after treatment with 20 µM FSK and 200 µM IBMX as denoted in the graph. The average fluorescence intensity of Cα in the nucleus was quantified in cells overexpressing the specified constructs.

The graph in Fig. 3D demonstrates the similar average fluorescent intensity of Cα in the nucleus when co-expressed with RIβ-WT, RIβ-L50R or RIβ-L50R-R211K. However, upon treatment designed to elevate cAMP, a robust increase in Cα fluorescence intensity was measured upon RIβ-L50R:Cα dissociation, as compared to that obtained upon RIβ-WT:Cα dissociation. Introducing the R211K replacement into RIβ-L50R further demonstrates the dependency of Cα translocation into the nucleus upon RIβ-L50R:C dissociation. This live cell experiment is consistent with the images of fixed cells captured before and after 30 and 60 min of cAMP level-affecting treatment (Fig.3A-C), in that both approaches demonstrated increased and rapid translocation of Cα into the nucleus upon elevation of cAMP levels in cells expressing the RIβ-L50R mutant.

### RIβ-L50R:Cα demonstrates faster holoenzyme dynamics in response to cellular cAMP signals than does the RIβ:Cα complex

To further investigate RIβ-L50R:Cα holoenzyme dynamics and to compare this heterodimer to the RIβ:Cα tetramer, we conducted a bioluminescence resonance energy transfer (BRET) assay. In this assay, the R and C subunits were fused with RLuc8 and GFP^2^, respectively, as illustrated in Fig. 4A. To this end, cells were co-transfected to co-express GFP^2^-labeled Cα subunit together with RLuc8-labeled RIβ-WT, RIβ-L50R or RIβ-G201E/G325E. The RIβ-G201E/G325E was unable to bind cAMP, and thus served as a negative control for cAMP sensitivity upon holoenzyme assembly (27, 28). First, cells overexpressing these vectors were treated with FSK and IBMX, resulting in increased intracellular cAMP levels. Cells over-expressing the RIβ-WT:Cα complex dissociated upon FSK/IBMX treatment whereas cells over-expressing the RIβ-G201E/G325E:Cα complex were unable to respond to the cAMP-affecting stimulus, and as such, this holoenzyme did not dissociate. Notably, the BRET curve of the RIβ-L50R:Cα complex resembled that of RIβ-WT:Cα, indicating that holoenzyme dissociation in the presence of the RIβ-L50R variant was similar to that of the native complex upon such stimulation (Fig. 4B, Fig. S1A). However, when exposed to 1 μM isoproterenol (ISO), a condition that generates relatively physiological cAMP concentrations (29–31), a faster heterodimer dissociation was observed in the presence of the RIβ-L50R variant than of RIβ-WT, as reflected by a more rapid decline in the BRET signal (Fig 4C, Fig. S1B). This decrease was followed by a rapid re-gaining of the BRET signal, resulting from a phosphodiesterase-dependent cAMP degradation, indicating a faster holoenzyme reassociation of the RIβ-L50R variant. These results suggest that the RIβ-L50R mutant compromises the precise regulation of the heterodimer dynamics.

**Figure 4.**
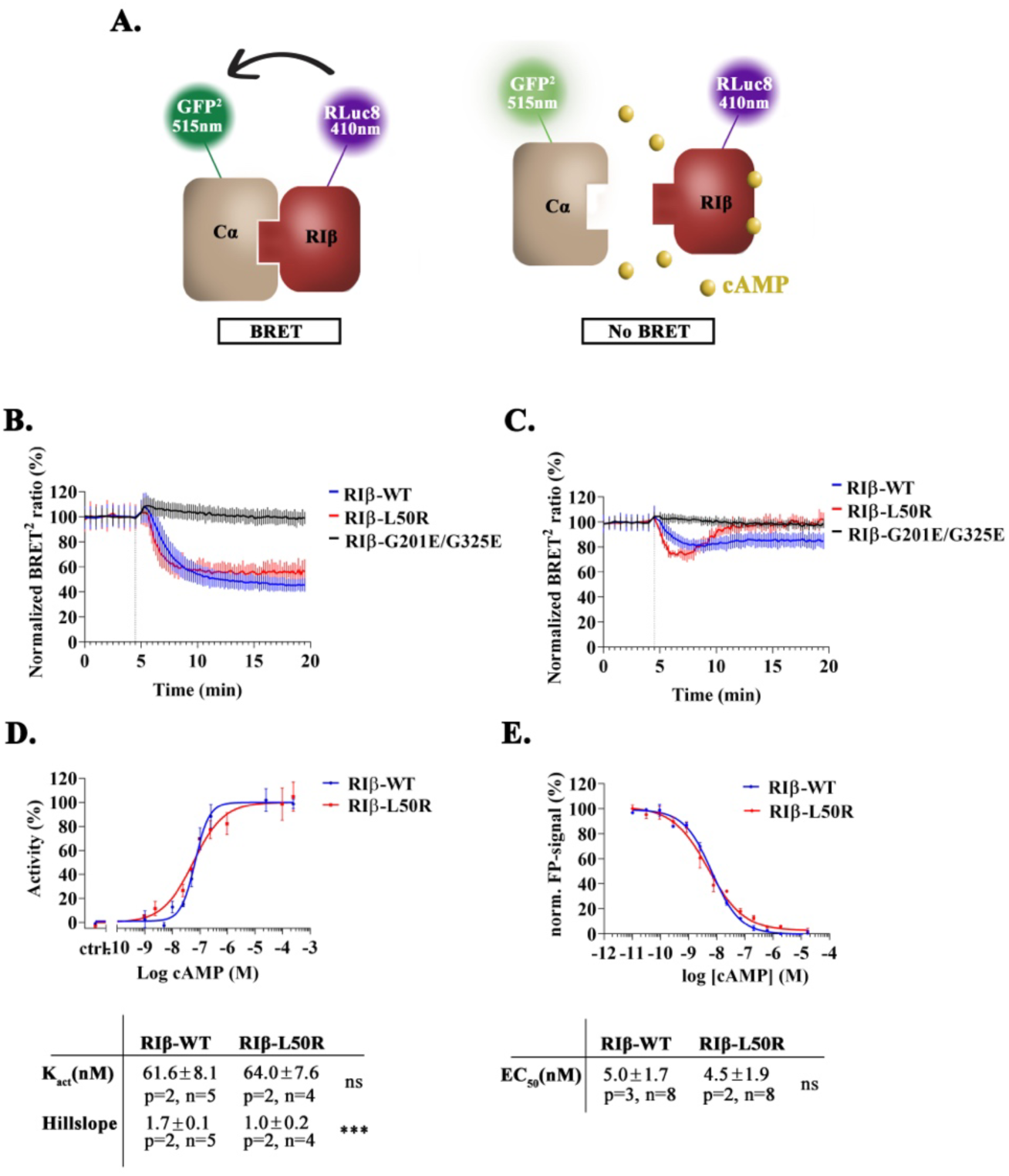
PKA holoenzyme dynamics are altered by the RIβ-L50R variant under cellular stimulation. **A.** Illustration of PKA holoenzyme dynamics analyzed by the BRET^2^ system. GFP^2^-tagged Cα and RLuc8-tagged RIβ are denoted. cAMP is represented by yellow dots. **B-C.** BRET signal profiles following stimulation of HEK293 cells with either 50 µM FSK and 100 µM IBMX **(B)** or 1µM ISO **(C)**. The BRET signal was monitored for 20 min. Data represent the normalized BRET^2^ ratio corresponding to the GFP^2^-Cα signal (515 nm) over a RIβ-RLuc8 signal (410 nm). BRET^2^ ratios were normalized between the Rluc8 signal (see Fig. S1 for raw data) and values measured before injection. RIβ-G201E/G325E served as a control for cAMP sensitivity. Each curve represents the mean of six replicates ± SEM. **D.** cAMP-mediated PKA holoenzyme activation as determined with a spectrophotometric kinase assay. Normalized values for PKA activity are plotted against the logarithmic cAMP concentration. Each measurement was performed in duplicate, and each curve represents the mean of the duplicates ± standard deviation (SD). Significance was tested for with an unpaired T-test (ns=non-significant, ***P≤0.001). Activation constants (K_act_) and Hill slopes were determined by applying a sigmoidal dose-response fits and are summarized in the table below (p=number of protein preparations, n=number of measurements). **E.** Fluorescence polarization assays measuring cAMP binding to RIβ-WT or RIβ-L50R. Statistical analysis showed no significance differences (ns: not significant, P> 0.05). All values are presented as the mean of multiple measurements ± SD, where “p” indicates the number of protein preparations and “n” indicates the number of measurements, each performed in duplicate. Significance was assessed using an unpaired t-test following confirmation of normal distribution.

To quantify the effect of RIβ-L50R substitution on PKA holoenzyme activation, we calculated the apparent activation constants (K_act_) for RIβ-WT:Cα and RIβ-L50R:Cα holoenzymes. The K_act_ for the RIβ-L50R variant was calculated *in vitro* and was identical to RIβ-WT holoenzyme (Fig. 4D). A substantial difference was, however, noted in the significantly lower Hill slope value calculated for the RIβ-L50R holoenzyme, pointing to a loss of cooperativity in cAMP binding upon holoenzyme activation (Fig. 4D). While the Hill slope value for the RIβ-WT:Cα holoenzyme indicated positive cooperativity, similar to a previously reported value for the isoform-specific RIα:Cα holoenzyme (32), the RIβ-L50R-containing holoenzyme exhibited Hill slope values comparable to those of a monomeric RIα deletion mutant (32). These findings further support the monomeric nature of RIβ-L50R and emphasizes the critical role of the D/D domain in promoting cooperativity. To explore whether the L50R variant affects cAMP binding to the RIβ subunit, we conducted a fluorescence polarization (FP) competition assay that measured the interaction between labeled cAMP and RIβ-WT or RIβ-L50R. As indicated in Fig. 4E, the EC_50_ values for cAMP were consistently in the low nanomolar range for both RIβ variants, reflecting a lack of significant difference and implying that the L50R variant has no effect on the affinity of the RIβ subunit to cAMP, as compared to the WT protein.

### Purified recombinant RIβ-L50R does not dimerize and is prone to aggregation

To complement the cellular studies described above and to directly evaluate the ability of RIβ-L50R to dimerize, we purified both full-length RIβ-WT and RIβ-L50R and subjected them to gel filtration chromatography (Fig. 5A-B). RIβ-WT eluted as in a single peak behaving as a stable dimer in solution, with a Stokes’ radius of 45.3 Å. In contrast, RIβ-L50R exhibited a distinct chromatographic profile containing multiple peaks, none of which corresponded to the expected size of a dimer (Fig. 5B). The significantly larger Stokes’ radius of peak I (63.1 Å), as compared to the expected value associated with the dimer, suggests the presence of higher-order multimers of this mutant RIβ protein (Fig. 5C). To further investigate the composition of the separated RIβ species, we performed Western blot analysis on the fractions collected (Fig. 5D). Fractions corresponding to peak II contained two distinct RIβ species. The first, appearing as a shoulder of the peak, corresponded to the expected monomeric RIβ protein, with a Stokes’ radius of 30.6 Å. A second species, also detected by the RIβ-specific antibodies, represents a truncated version of the RIβ subunit (25.7 Å), likely resulting from proteolytic degradation. The Stokes’ radius of this truncated species is similar to that of the RIα monomeric deletion mutant (RIα-Δ1-91) that does not include the sequence corresponding to the D/D domain (32), suggesting the instability and proteolytic susceptibility of RIβ-L50R. These data support the conclusion that in the RIβ-L50R variant not only dimer formation is disrupted but also protein stability is weakened and promotes aggregation.

**Figure 5.**
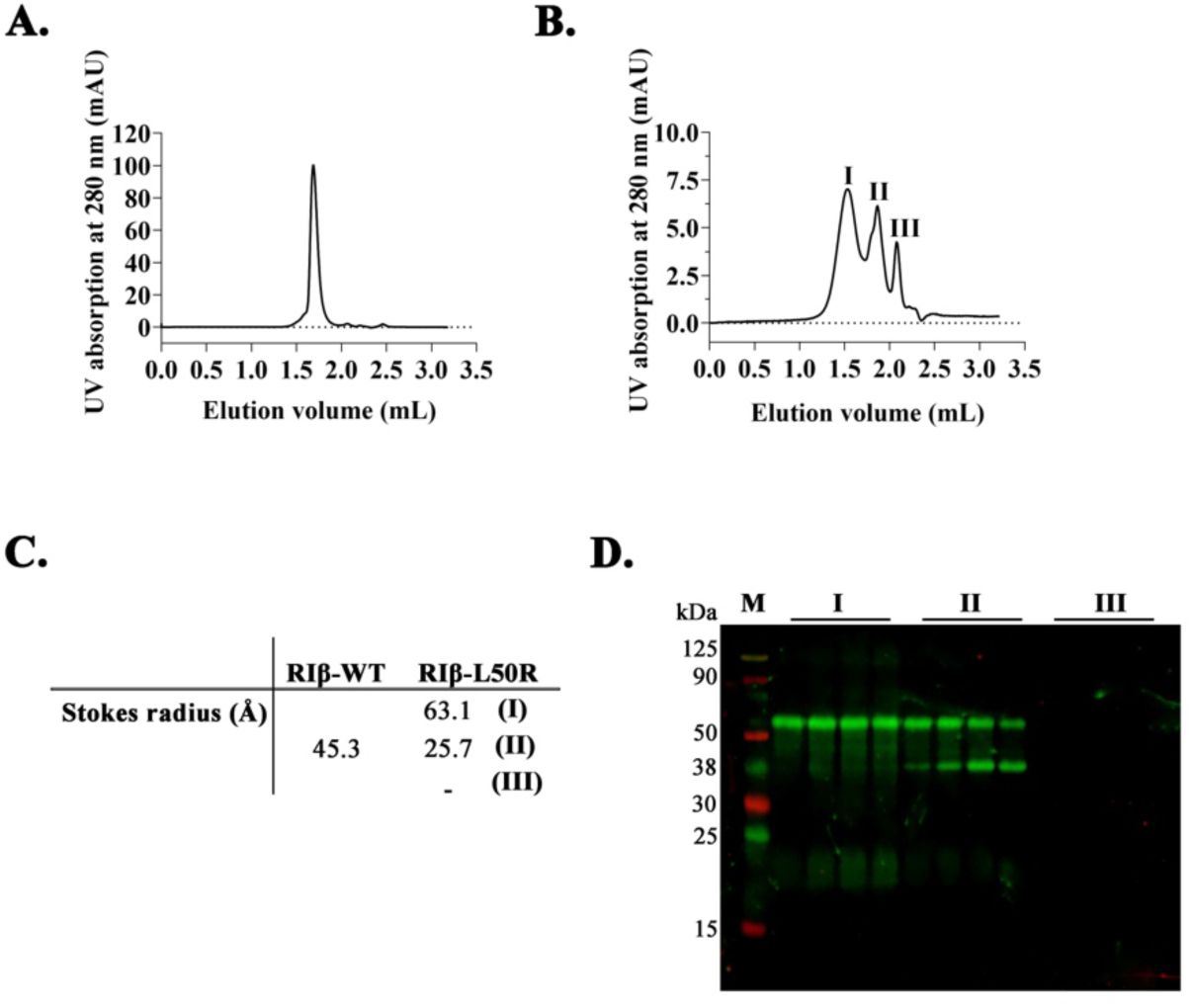
Disrupted dimerization and aggregation tendency of purified RIβ-L50R proteins. **A-B** Size exclusion chromatography profiles of **(A)** RIβ-WT and **(B)** RIβ-L50R. **C.** Calculated Stokes radii derived from the elution volumes of indicated peaks for each purified protein. **D.** Western blot analysis of fractions from peak I, II, and III of the RIβ-L50R purified protein using RIβ-specific antibodies.

### The RIβ-L50R variant shows enhanced enhances cAMP-induced dissociation of the RIβ-L50R:Cα holoenzyme

Surface plasmon resonance (SPR) measurements were conducted to assess the impact of the L50R replacement on the RIβ:Cα affinity. FSS-Cα was captured on a streptactin coated sensor chip and RIβ-WT or RIβ-L50R, respectively, were injected in the presence of MgATP to determine the affinities between the respective R subunits and the C subunit. The calculated K_D_ values for the RIβ-WT:Cα and RIβ-L50R:Cα interactions were 0.53 ± 0.34 nM and 0.17 ± 0.14 nM, respectively, indicating sub-nanomolar affinities between RIβ-WT or RIβ-L50R and the Cα subunit (Fig. 6A-C). These strong affinities are based on rapid association and slow dissociation phases (Fig. 6A-B).

**Figure 6.**
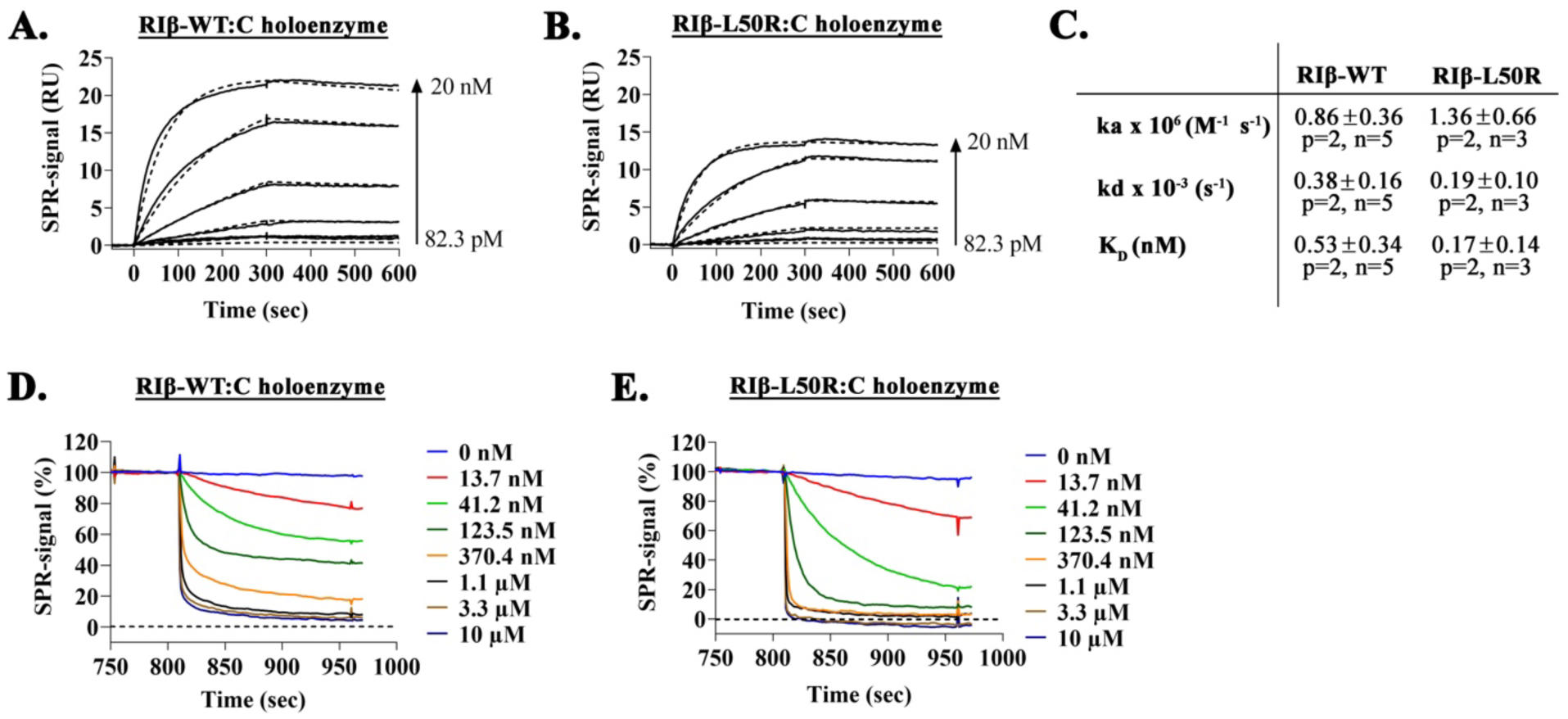
The RIβ-L50R:Cα holoenzyme requires less cAMP to dissociate, despite high affinity binding to the Cα. **A-B.** Binding kinetics of RIβ-WT **(A)** and RIβ-L50R **(B)** to immobilized FSS-Cα subunit measured by SPR. **C** Kinetic and equilibrium binding constants, determined using a 1:1 Langmuir binding model are summarized. Values are the mean of multiple measurements ± SD; p = number of protein preparation, n = number of measurements. Data were evaluated with Biacore T200 evaluation software 3.2 **D-E.** cAMP-induced dissociation of RIβ subunits from preformed holoenzymes was determined by SPR. Increasing cAMP concentrations were injected during the dissociation phase for RIβ-WT:C **(D)** and RIβ-L50R:C **(E).** Sensograms represent changes in the normalized SPR signal over time. Original sensograms are represented in Figure S2. RU: response units.

To explore the effect of cAMP on the dissociation of R subunits from the immobilized Cα subunit, increasing concentrations of cAMP were added during the dissociation phase of the SPR assay (Fig. 6D-E and Fig. S2). RIβ-L50R dissociated at lower cAMP concentrations than those required by RIβ-WT, suggesting a less stable mutant holoenzyme (Fig. 6D-E).

### Transcriptome analysis of patient primary fibroblasts

The biochemical and cellular data showing that PKA allosteric regulation is affected by the mutation resulting in RIβ-L50R prompted us to perform transcriptome analysis so as to comprehensively understand the molecular consequences of unregulated PKA and its correlation with clinical symptoms. Analysis of the transcriptome of primary fibroblasts from a female patient expressing the RIβ-L50R variant and diagnosed with a neurodegenerative disorder, relative the transcriptome of fibroblasts from a healthy age- and sex-matched individual, revealed substantial alterations in gene expression (Fig. 7). Of a total of 17, 833 genes identified, with 9, 002 were up-regulated and 8, 831 were down-regulated (ShinyGO v0.741). Only genes exhibiting fold changes <-2 or >2, which were statistically significant at padj<0.05, in the patient relative to the control were further analyzed (Fig. 7). Disease-based classification of the transcriptome revealed a gene expression signature associated with Alzheimer’s disease (Fig. 7C1), as reflected by changes in the levels of transcripts for cellular components such as neuron projections, synapses, axons, and dendrites (Fig. 7C2). Notably, genes involved in locomotion function were significantly changed in the patient expressing the RIβ-L50R mutant, consistent with the patient’s clinical symptoms. Furthermore, we observed differential expression of key hub genes implicated in neuronal generation, neuronal survival and growth, which correlated with MRI scans indicating pronounced brain atrophy, surpassing what is typical at the patient’s age. Interestingly, the patient was prescriped of medication for hypertension. This aligned with the differential expression in a notable gene cluster associated with this condition, furthur validating our analysis.

**Figure 7.**
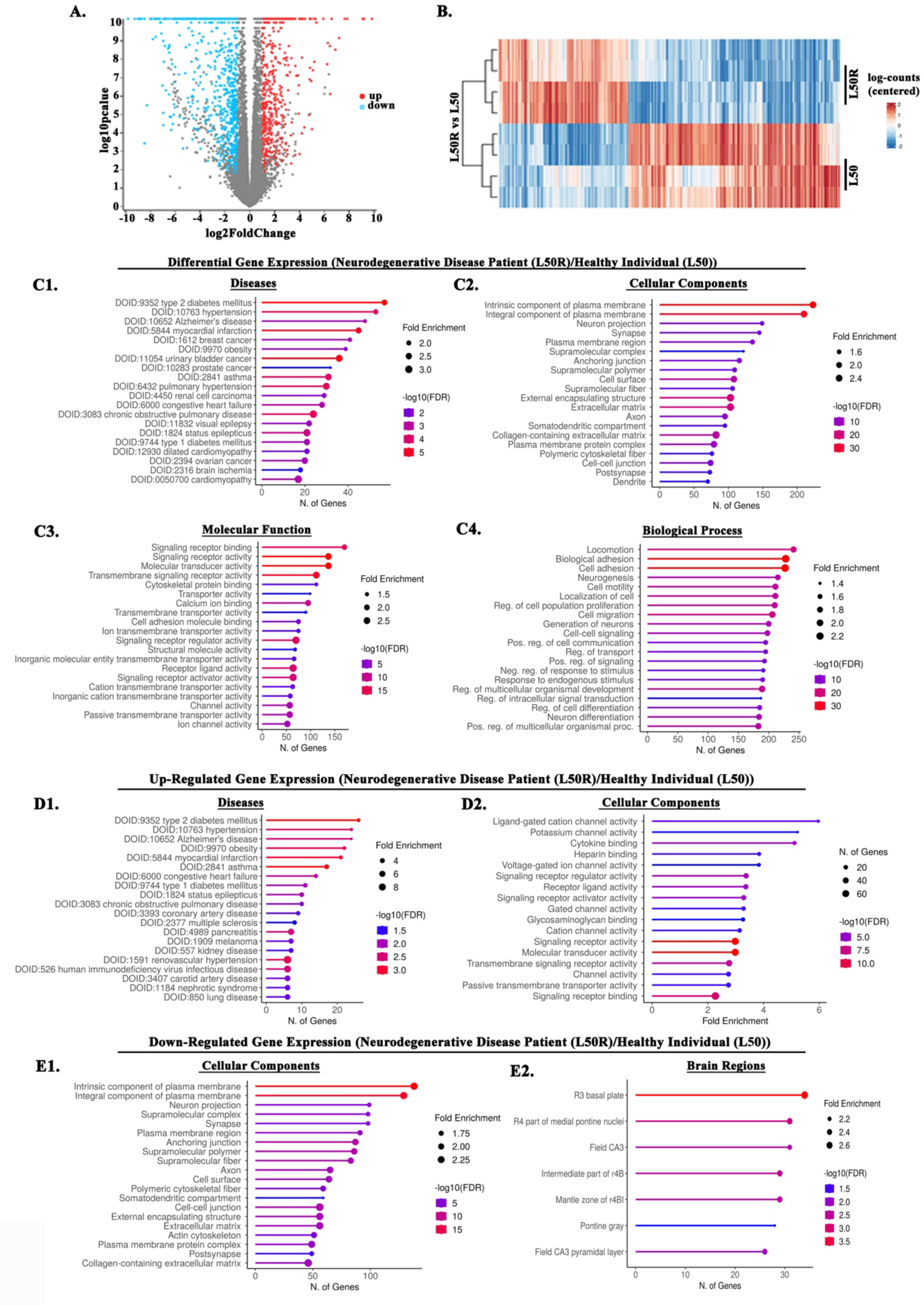
Differential gene expression analysis of primary human fibroblast cells. Analysis of the transcriptome changes in primary fibroblasts comparing a female patient expressing the RIβ-L50R variant, diagnosed with a neurodegenerative disorder, and that of a healthy individual. Analysis was conducted using ShinyGO: Gene Ontology Enrichment Analysis. **A.** Volcano plot displaying differential expression. Blue color represents down-regulated genes while red color represents up-regulates genes in affected individual (RIβ-L50R), as compered to the healthy individual (RIβ-L50). Genes with a log_2_bfold change ≥ 1 (corresponding to a multiple hypothesis-adjusted p-value (padj) ≤0.05) are highlighted. **B.** Heatmap illustrating gene expression patterns. Blue shading indicates down-regulated genes, and red shading indicates up-regulated genes in the RIβ L50R variant-expressing subject, as compared to the RIβ-L50 variant-expressing subjuect. Darker shading indicates higher significance. **C.** Differentially expressed genes that are up- and down-regulated in the RIβ L50R variant-expressing subject, as compared to the RIβ-L50 variant-expressing subject. Pathway classification analysis as denoted (C1-4). **D.** Pathway classification analysis of up-regulated genes in the RIβ L50R variant-expressing subject, as compared to the RIβ-L50 variant-expressing subject (D1-2). **E.** Pathway classification analysis of down-regulated genes in the RIβ L50R variant-expressing subject, as compared to the RIβ-L50 variant-expressing subject (E1-2). The false discovery rate wass calculated based on the nominal p-value from the hypergeometric test. Fold-enrichment is defined as the percentage of genes in a list belonging to a pathway, divided by the corresponding percentage in the background.

## Discussion

In this study, we uncovered the molecular mechanisms underlying the clinical manifestations of neurodegeneration in patients expressing the RIβ-L50R variant, revealing disrupted PKA holoenzyme assembly and uncontrolled allosteric regulation. Utilizing patient-derived cells, as well as direct measurements of purified proteins, we shed light on the critical position of RIβ L50 in facilitating the dimer formation of the regulatory subunit, crucial for the precise regulation dynamics of the PKA holoenzyme.

Exploration of *PRKARIB* variants via database screening uncovered a clustering of mutations predominantly within that region encoding cAMP-binding sites, with very few mutations observed in that region encoding the D/D domain. This pattern suggests that the D/D domain is evolutionary conserved, emphasizing its essential function in maintaining structural integrity and allostery of holoenzyme assembly.

Our focused analysis on L50R PRKARIB variant, the only known PKA regulatory subunit variant confirmed to disrupt dimer formation, not only unraveled dynamics properties of PKA assembly and allosteric regulation but also provided insights into the functional consequences leading to neurodegenerative disease. Although other pathogenic RIβ variants, such as the I40V, R68Q and A67V mutants, presented replacements of residues localized within the D/D domain, they did not disrupt dimerization and thus were not the primary focus of our study.

Integration of structural analysis results with those obtained in biochemical studies employing purified proteins, and cellular studies involving cells over-expressing targeted fluorescent proteins, as well as patient-derived cells, consistently indicated that the mutation leading to the appearance of the RIβ-L50R variant disrupted dimer formation and rendered the protein prone to aggregation. Gel filtration analysis revealed distinct differences between recombinant RIβ-WT and RIβ-L50R proteins, with RIβ-WT exhibiting a stable dimeric structure while RIβ-L50R displaying a high tendency to form multi-oligomers. CD spectroscopy analysis corroborated these findings, demonstrating alteration in secondary structure and stability of the protein, in line with predictions from the *in-silico* modeling.

In cells, RIβ-L50R aggregates were detected in the insoluble fraction. Relying on the approaches listed above, we were able to conclude that such aggregation resulted from structural perturbation and misfolding of the D/D domain. Consequently, the docking site for AKAPs within this domain failed to appear, leading to the mutant protein being unable to bind dAKAP1. This emphasizes the critical importance of proper folding of this PKA R subunit domain for assembly of macromolecular complexes.

The molecular mechanism elucidated here for a PKA dependent neurodegenerative disease in which protein aggregation results from disrupted homodimerization, sheds light on a potentially common but under-appreciated mechanism shared by several neurodegenerative diseases. Mutations in the *DJ-1* gene associated with autosomal recessive early-onset Parkinsonism results in destabilized homodimerization (33, 34). The crystal structure of human DJ-1 demonstrated the protein to exist as a stable homodimer. The mutation leading to L166P replacement in DJ-1, identified as causal in patients with Parkinson’s disease, produces an unstable form of the protein that forms higher-order oligomers and is degraded by the proteasome (35). Similarly, mutations in SOD1 associated with amyotrophic lateral sclerosis (ALS) share a similar molecular mechanism of dimer destabilization (36–38). This emphases the broader significance of disrupted homodimerization in protein aggregation and disease pathology.

We previously solved the crystal structure of the RIβ holoenzyme as a dimer of two heterodimers composed of a regulatory and catalytic subunit (39). In the absence of a folded dimerization domain, RIβ-L50R can interact with the catalytic subunit to form a heterodimer instead of a dimer of heterodimers (Fig. 8). We have shown that interaction of RIβ-L50R with a catalytic subunit protects the former from aggregation (25). In the present study, we sought to dissect the allosteric network of the RIβ holoenzyme and determine how the mutation leading to expression of the L50R variant impacts PKA holoenzyme dynamics. We showed that RIβ-L50R binds with high affinity to the C subunit, as measured by SPR. The retained high affinity of the RIα regulatory subunit that lacks the N-terminal D/D domain (RIα-Δ1-91) for the C subunit was shown earlier (32). Our results confirmed that the RIβ:Cα interactions for one heterodimer are remarkably similar for the different RI isoforms (16), and that the differences between the isoforms occur at the tetrameric level. The presence of RIβ-L50R in its monomeric, rather than its dimeric form led us to investigate the subcellular localization of RIβ-L50R and Cα in primary fibroblasts from both a healthy individual and a patient with the mutation. While the localization of RIβ remained unchanged in the presence of the mutation, an increase in Cα expression within the nucleus was evident in the patient expressing the mutant protein. This increase was also replicated in PC12 cells treated with cAMP stimulators, suggesting that the RIβ-L50R:C heterodimer dissociates more easier than does the RIβ_2_:C_2_ holoenzyme in the presence of cAMP within the cell. This is in line with the BRET results reported here.

**Figure 8:**
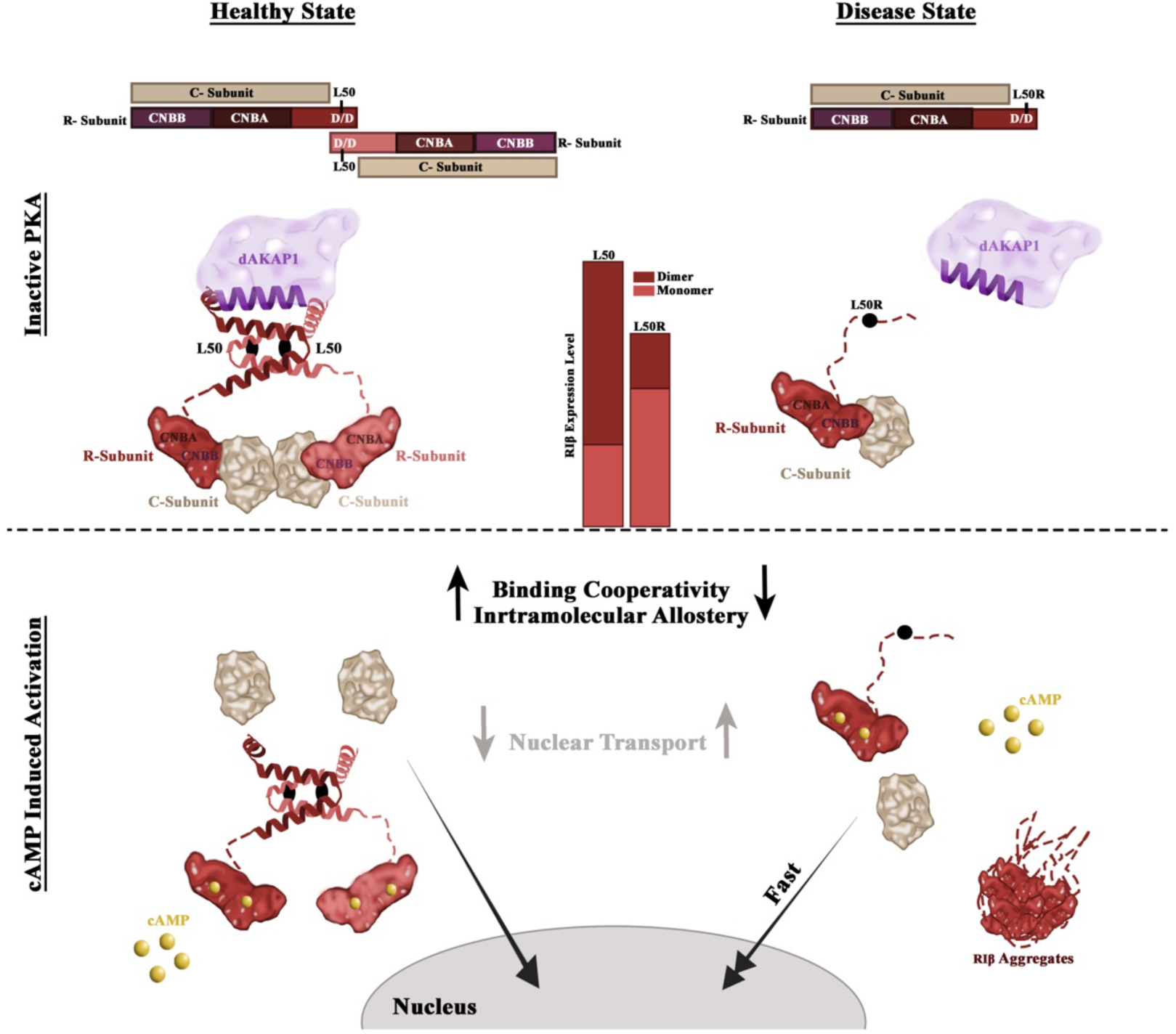
Illustration of PKA holoenzyme assembly and dynamics in the healthy and disease state before and in response to cAMP stimulation. Upper panel depicting the inactive state. The upper panel depicts the inactive state. In the healthy state, the PKA holoenzyme is a dimer of two RIβ:C heterodimers. Homodimerization provides a binding site for AKAPs. In the disease state, PKA exists as a RIβ:C heterodimer, where the L50R mutant perturbs homodimerization and AKAP binding. The bars in the illustration represents RIβ expression levels and the ratio between monomer and dimers. The lower panel depicts PKA dynamics upon cAMP-induced activation. In the healthy state, the RIβ dimer dissociates from the C subunit, allowing its translocation into the nucleus. In the disease state, the RIβ-L50R:C heterodimer dissociates faster, exhibits reduced cooperativity, leads to RIβ aggregation and promotes faster translocation of the C subunit into the nucleus.

Our results further demonstrated that the presence of the L50R substitution resulted in a RIβ-L50R:C heterodimer complex that exhibited increased sensitivity to cAMP, requiring lower concentrations for dissociation than did the complex containing the native RIβ protein. This mutant-incorporating heterodimer displayed reduced cooperativity, as compared to the native complex, as evident from the Hill slope values. These results emphasize the role of the RIβ N-terminal domain in maintaining cooperativity of the PKA holoenzyme and suggest that the L50R mutant compromises the precise regulation of heterodimer dynamics. Based on the findings of this study, we propose that decreasing the affinity of RIβ-L50R to cAMP could offer a promising strategy to restore regulation over the reduced allosteric mechanisms caused by the associated mutation. This approach thus presents a potential avenue for therapeutic intervention.

Our thoroughly comprehensive investigation of the dynamics of the RIβ-L50R:C heterodimer, ranging from *in vitro* analysis of the purified proteins to visualization of complex assembly and dynamics in cells derived from a patient affected by a neurodegenerative disease, helped unraveled the intricate interplay between PKA holoenzyme structure in the presence of the mutant, disrupted assembly and dynamics, and the onset of neurodegenerative phenotypes. These phenotypes are reflected by a unique pattern of gene expression specific to the disease (i.e., NLPD-PKA), which shares commonalities with changes observed in other neurodegenerative diseases. For instance, alternations in the expression of key hub genes associated with neuronal degeneration, growth, and survival are indicative of the severe brain atrophy observed in MRI scans of affected patients expressing the RIβ-L50R variant. This pattern of gene dysregulation mirrors findings seen in other neurodegenerative diseases, such as Alzheimer’s disease, where similar alternations in gene expression profiles have been reported (40). Furthermore, the observed dysregulation of cellular components, including neuron projections, synapses, axons and dendrites in the patient expressing the RIβ-L50R variant is consistent with the pathological changes seen in other neurodegenerative diseases. Examples of such diseases include Parkinson’s disease, where alternation in neuronal projections and synapses contribute to motor impairments (41), Huntington’s disease, characterized by abnormalities in axonal transport and dendritic spine densities resulting in cognitive dysfunction (42), and ALS, where abnormalities in axonal transport and axonopathy contributes to progressive muscle weakness (43)

These changes reflect synaptic disruption and neuronal loss, as well as disrupted neuronal circuits, all of which are common features of observed in various neurodegenerative conditions. Given that PKA dysfunction contributes to the pathogenesis of various neurodegenerative diseases (8–13), establishing a direct link between unregulated PKA allostery and assembly and the products of genes associated with neurodegeneration will help identify therapeutic targets for maintaining PKA function.

## Acknowledgments

We thank all members of the Ilouz lab for helpful discussions. We thank Dr. Shai Bel for helpful discussions on RNA sequencing data. We thank M. Hansch, S. Kasten and O. Bertinetti of the Herberg lab for excellent technical assistance. We would also like to thank Choel Kim (Baylor College of Medicine, Houston, United States) for providing the *E. coli* TP2000 cells, Sabrina Dörfler and Maximilian Wallbott (University of Kassel, Department of Biochemistry, Germany) for the pQTEV RIβ plasmids used for the recombinant protein expression. RI was supported by the Israel Science Foundation (grant number 1706/22) and by the US-Israel Binational Science Foundation (2019270). FWH acknowledges the financial support by the German Research Foundation (DFG)-funded Research Training Group “multiscale clocks” (448909517/GRK 2749: Biological Clocks on Multiple Time Scales).

## Material And Methods

### Subjects

Primary human fibroblast cells were obtained from the Erasmus Medical Center in Rotterdam, Netherlands. Informed consent was obtained from all participates or their legal representatives. Age- and sex-matched control subjects exhibited no neurodegenerative changes upon pathological examination. The study recived ethical approval under number: 2009-401.

### Cell culture

Primary human fibroblast cells, rat pheochromocytoma (PC12) cells, and human embryonic kidney 293 (HEK293) cells were maintained in high glucose Dulbecco’s modified Eagle’s medium (DMEM) (Biological Industries) supplemented with 10% fetal bovine serum (Biological Industries), 2 mM L-glutamine (Sartorius) and 5% penicillin-streptomycin (Biological Industries) at 37°C in an atmosphere of 5% CO_2_. All cell lines were negative-mycoplasma.

### Mutations

Single-site mutations that led to the replacement of the indicated amino acids at the following position I40V, L50R, A67V, R68Q, and R211K were introduced into the mKO2-RIβ plasmid using a site-directed mutagenesis kit (New England BioLabs, Ipswich, MA). N-terminal His-tagged human RIβ-WT and RIβ-L50R constructs, cloned in pQTEV plasmids, were used to express recombinant RIβ proteins for *in vitro* biochemical analysis. All constructs confirmed by sequencing.

### Antibodies

Primary antibodies: Sheep anti-PKA RIβ antibodies (R&D Systems catalog # AF4177 (RRID: AB_2284184)) were diluted 1:2, 500 for Western-Blot (WB), and 1:100 for immunohistochemistry (IHC). Mouse anti-PKAc monoclonal antibodies BD Biosciences catalog # 610981 (RRID: AB_398294) were diluted 1:4, 000 for WB, 1:50 for IHC. Rabbit anti-GAPDH Abcam Catalog # ab9485 (RRID: AB_307275) dilution 1:2, 500 for WB. Rabbit anti-Topoisomerase I Abcam Catalog # ab109374 (RRID: AB_10861978) dilution 1:1, 000 for WB. Ranit Anti-AKAP1 Cell signaling Catalog #CST-5203 (RRID: AB_10828202) dilution 1:1, 000 for WB, 1:100 for IHC. The WB primary antibodies were prepared in 1% filtered BSA in PBST solution, and the IHC primary antibodies were prepared in 1:10 blocking solution (10% Normal Donkey Serum (Jackson ImmunoResearch) + 0.1% filtered BSA) in PBST. Rabbit Anti-PKA-RIβ (ABM catalog #Y051648) diluted 1:2, 000 in TBS-T buffer (20 mM Tris, pH 7.5, 140 mM NaCl and 0.1% (v/v) Tween 20) supplemented with 1-2% (w/v) milk powder was used as primary antibody for the WB of the analytical gel filtration fractions. Secondary antibodies: Donkey Anti-Sheep IgG, HRP Conjugated (Abcam) (RRID: AB_955452), Goat Anti-Rabbit IgG, HRP Conjugated (Abcam) (RRID: AB_955447), Rabbit to Mouse IgG, HRP Conjugated (Abcam) (RRID: AB_955440), Donkey Anti-Sheep IgG (Alexa Fluor 647) Abcam Catalog # ab150179, Goat Anti-Rabbit IgG (L+H) (Alexa Fluor 488) Invitrogen Catalog # A-11934. All HRP-secondary antibodies were used at 1:10, 000 dilution and all fluorescent-secondary antibodies were used at 1:250 dilution. IRDye 800CW Donkey Anti-Rabbit (LI-COR Biosciences) (catalog # 926-32213) diluted 1:15, 000 in TBS-T buffer with 1-2% (w/v) milk powder was used as a secondary antibody for the WB of the analytical gel filtration fractions.

### Transient transfection

PC12 cells were grown into 6-well plates for WB and in a 24-well plate for immunofluorecence (IF). After 24 h of incubation the cells were transfected with 2M Calcium Chloride and DNA (0.5 µg for 24 well or 5 µg of 6 well plates). The solution was added to HEPES-buffered saline solution with the same volume and incubated for 15 min at room temperature. After incubation, the transfection solution was gently added to the cell media gently, and the cells were lysed for WB or fixed for IF after 48 h.

For time-dependent BRET^2^ measurements 2x10^4^ HEK293 cells per well were seeded on a 96-well-microtiter plate and grown for 24h at 37°C at 6% CO_2_. After 24h 0.05 µg DNA of C-terminally RLuc8 (44) tagged WT or L50R RIβ-subunits and 0.05 µg DNA of N-terminally GFP^2^ tagged C-subunits per well were transfected. Cationic polyethyleneimine (PEI, Polysciences GmbH) solution was used to improve transfection. 24 h after transfection medium was exchanged with fresh DMEM medium including 10 % (v/v) 10% FBS (Capricorn Scientific).

### Cell fractionation and WB

Primary human fibroblasts were grown to approximately 70% confluence in 10 mm plates and incubated for 48 h or 3 X 10^5^ PC12 cells were grown in 6-well plate for 24 h, transfected and incubated for 48 h. Both cell types were washed twice in cold PBS (Biological Industries), suspended in lysis buffer containing 50 mM Tris-HCI pH 7.5, 0.1% Triton X-100, 150 mM NaCl, 1 mM EGTA, 1% NP-40, 0.25% sodium deoxcycholate, 1 mM PMSF and a cocktail of protease inhibitors (P8340 Sigma) (diluted 1:100) for primary fibroblasts or in lysis buffer containing 50 mM Tris-HCl pH7.4, 150 mM NaCl, 1 mM EDTA, 0.1% Triton X-100, 1% NP-40, 10 mM NaF, 1 mM Na_3_VO_4_, phosphatase inhibitors and protease inhibitors for PC12 cells, and incubated on ice for 15 min. The lysates were centrifuged at 15, 000 rpm and 4 °C for 30 min. for PC12 cells, after centrifugation, the supernatant was collected as soluble fraction and the pellet was washed twice in cold PBS, resuspended in lysis buffer containing 6 M urea, and sonicated. Sample buffer X5 with or without SDS and β-mercamtoethanol (for non-reduced conditions) was added, and the samples with reduced conditions were boiled for 4 min. Protein lysates were resolved by SDS-PAGE and transferred to PVDF membranes (Bio-Rad) using a Trans Blot Turbo RTA midi transfer kit (Bio-Rad). The PVDF membranes were blocked with 5% Bovine serum albumin (BSA) in PBST (0.1% Tween 20) for 1 h at room temperature, and subsequently incubated overnight at 4 °C with primary antibodies. After incubation, the PVDF membranes were washed four times with PBST for 10 min. secondary antibodies with HRP were incubated for 1 h. After four washes with PBST, the PVDF membranes were detected using ECL.

### Immunohistochemistry and Immunofluorescence

Primary fibroblasts were grown on 13 mm coverslips in 24-well plates to approximately 70% confluence and incubated for 48 h. After treatment the cells were washed twice in PBS and fixed for 15 min in 4% paraformaldehyde. After fixation, cells were washed five times with PBS and permeabilized and blocked in PBS containing 0.5% Triton-X100 and 1% BSA for 30 min. After blocking, cells were immunostained for overnight at 4 °C with primary antibodies. After this time, the cells washed three times in PBS and incubated with a secondary antibody for 1 h at room temperature and washed twice with PSB. Nuclei were counterstained with Hoechst (Invitrogen) for 5 min. For PC12 cells, 0.6 X 10^5^ PC12 cells were grown on 13 mm coverslip in a 24-well plate for 24 h, co-transfected and incubated for 48 h. After treatment the cells were washed twice in PBS and fixed for 15 min in 4% paraformaldehyde then washed three times with PBS and stained with Hoechst for 5 min. The slides mounted on cover glass with Gelvatol and were imaged on a Zeiss LSM 780 confocal microscope and Leica STED live imaging microscope.

### PKA RIβ Dimerization/Docking Domain Expression and Purification

Wild-type PKA RIβ Dimerization/Docking (D/D) domain was expressed in *Escherichia coli* BL21 (DE3) pLysS at 37°C until an OD of 0.6 was reached. IPTG was added in a final concentration of 0.5mM at 18°C overnight. Cells were harvested by centrifugation at 5, 000 g for 11 minutes at 4°C and then resuspended with a ratio of 1:10 w/v Lysis Buffer (50 mM Tris, 500 mM NaCl, 1 mM DTT, pH 7.5) supplemented with protease inhibitors (10 mM Benzamidine, 0.4 mM 4-2-Aminoethyl-benzenesulfonyl fluoride, 1 mM Pepstatin, 1 mM Leupeptin, 28 mM Tosyl-phenylalanyl-chloromethyl-ketone and 28 mM Tosyl-L-lysylchloromethane-ketone). Cells were then lysed with a Microfluidizer at 10, 000 psi. The homogenized mixture was separated by centrifugation at 15, 000 g for 45 minutes at 4°C. The supernatant was treated with 45% ammonium sulfate for 1 hour at 4°C to precipitate the soluble proteins. The precipitate was collected via centrifugation at 10, 000 g for 10 minutes at 4°C, then solubilized with Lysis Buffer followed by another round of centrifugation at 16, 000 g for 10 minutes at 4°C. The supernatant was then added to a nickel-coupled agarose resin and incubated overnight at 4°C. The nickel resin had been equilibrated with Lysis Buffer prior to incubation with the supernatant. The next day, protein was eluted from the resin using Elution Buffer (50 mM Tris, 500 mM NaCl, 500 mM imidazole, pH 7.5). The eluted protein was unfolded inf 8M urea with 5 mM DTT, and incubated for 30 minutes at room temperature. Step-wise dialysis using Gel Filtration Buffer (50 mM Tris, 200 mM NaCl, pH 7.5) at 4°C was performed over the next 24 hours to remove the urea and DTT. The refolded protein was separated by size-exclusion chromatography (HiLoad 16/600 Superdex 200 pg, Cytiva) via FPLC (NGC, Bio-Rad) to remove protein aggregates. Protein from the peak corresponding to the expected dimer size, ∼90mL, was collected, combined and reinjected into the size-exclusion chromatography to demonstrate the protein would remain stable as a dimer and not reform aggregates. The purified protein was stored in Gel Filtration Buffer + 25% (v/v) glycerol at -80°C.

The L50R mutant was expressed, harvested and lysed like wild-type. However, L50R expressed mainly in inclusion bodies, which were extracted at room temperature using a ratio of 1:2 w/v Extraction Buffer (8M urea, 50 mM Tris, 500 mM NaCl, 5 mM DTT, pH 7.5 + the aforementioned protease inhibitor cocktail). After extraction, the solution was centrifuged at 15, 000 g for 15 minutes. The supernatant underwent refolding via step-wise dialysis in Gel Filtration Buffer at 4°C over the next 24 hours to remove the urea and DTT. After refolding, the suspension was mixed with nickel-coupled agarose resin and incubated overnight at 4°C. The nickel resin had been equilibrated with Gel Filtration Buffer prior to incubation with the supernatant. The next day, protein was eluted from the resin using Elution Buffer (50 mM Tris, 200 mM NaCl, 500 mM imidazole, pH 7.5). Protein was then separated by size-exclusion chromatography (HiLoad 16/600 Superdex 200 pg, Cytiva) via FPLC (Bio-Rad). Purified protein was stored in Gel Filtration Buffer + 25% (v/v) glycerol at -80°C.

### Secondary Structure Analysis of the PKA RIβ D/D Domain

Circular dichroism (CD) was used to evaluate the secondary structure D/D domain of PKA RIβ wild type and L50R. Experiments were performed using a Jasco J-715 spectrometer and all signals were corrected for buffer effects (50 mM Tris, 200 mM NaCl, pH 7.5. The data was collected using 15 μM of protein, with the following instrument parameters: continuous mode from 200-260 nm, pathlength = 0.1 cm, sampling rate 10 nm/min and a bandwidth of 10.0 nm. 13 and 9 scans were averaged for wild-type and L50R, respectively. Data analysis to estimate secondary structure was performed using BestSel (45–49).

### cAMP inducer experiment

48 h after PC12 cells or primary fibroblasts transfection, the media was replaced with serum-free DMEM and incubated for 1 hr.Then pharmacological agents were introduced for primary fibroblasts: 200 µM Isobutylmethylxanthine (IBMX, Sigma) and 1 µM isoproterenol (ISO, Abcam) that gently added to the cell media for 30 min and fixed to IF assay. PC12 cells treated with a cocktail pf 20 µM Forskolin (FSK, Abcam) and 200 µM IBMX (Sigma) and incubated for 30 min and 1 h and before fixation for IF assay.

### Live-cell imaging

3 X 10^5^ PC12 cells were grown on 35 mm glass bottom dish. After 24 h, the cells were co-transfected and allowed to incubate for 48 h. One hour prior the imaging, the medium was replaced with serum-free DMEM. The cells were imaged every 5 min for 20 min at five different positions. Then, the cells were treated with 20 µM FSK and 200 µM IBMX and imaging of the same positions continued at 5 min intervals. Images quantification was conducted using image J software.

### Image processing and relative quantification of laser intensity in the nucleus

A customized code written in Java for the Fiji (Version 1.54g 18 October 2023). Image processing platform was used to relatively quantify the intestisy of C-subunit in the neucleus. The Hoechst channel served to deliniate the nuclear region, with image aquired a magnification of 63X to enshure acccurate analysis.

### RNA extraction from primary fibroblast

Primary human Fibroblasts were washed with PBS before total RNA extraction using the RNeasy Plus Mini Kit (Qiagen).

### RNA sequencing and analysis

Integrity of the isolated RNA was tested using the Agilent High Sensitivity RNA Kit and Tapestation 4200 at the Genome Technology Center at the Faculty of Medicine Bar-Ilan University. 500 ng of total RNA was used for mRNA enrichment by using the NEBNext mRNA polyA Isolation Module(NEB, #E7490L) and libraries for Illumina sequencing were performed using the NEBNext Ultra II RNA kit (NEB, #E7770L). Quantification of the library was performed using dsDNA HS Assay Kit and Qubit 2.0 (Molecular Probes, Life Technologies), and qualification was done using the Agilent D1000 Tapestation Kit and Tapestation 4200. 400 nM of each library was pooled together and was diluted to 4 nM according to NextSeq manufacturer’s instructions. 1.6 pM was loaded onto the Flow Cell with 1% PhiX library control. Libraries were sequenced on an Illumina NextSeq 500 instrument, with 75 cycles of single read sequencing. Sequencing data was aligned and normalized at The Nancy and Stephen Grand Israel National Center for Personalized Medicine at the Weizmann Institute of Science. Pathways and Pathological analyses were performed using the ShinyGO web tool.

### Time-dependent BRET^2^ measurements

Time-dependent BRET^2^ measurements were performed at 37°C at the microplates reader Polarstar Omega device (BMG LABTECH). Emission from the RLuc8 (at 410 nm) and GFP^2^ (at 515 nm) signals was detected and ratio between the GFP^2^ and RLuc8 signal (BRET^2^-ratio) was calculated. For each construct, 6 wells were measured, and the mean values of the BRET^2^-ratio were plotted against time. In the first 4.5 minutes of the measurements, the signal without any stimulus was recorded. After 4.5 minutes of detection different cAMP stimuli were applied to the cells to induce holoenzyme dissociation. Holoenzyme dissociation was achieved using two different stimuli that increase the intracellular cAMP concentration. Cells were stimulated either with Isoproterenol (1 µM) or with a combination of Forskolin (50 µM) and the non-specific PDE inhibitor Isobuthylmethylxanthin (IBMX) (100 µM), to prevent degradation of cAMP in the system, and the resulting signal was detected in the following 15.5 minutes. Coelenterazin 400a (DeepBlueCTM, DBC, Biotrend) in a final concentration of 5 µM per well was used as substrate for the Luciferase enzymatic reaction. All dilutions for the cAMP stimuli and DBC were prepared in HBSS buffer (Hank’s Balanced Salt Solution w/o Mg^2+^/Ca^2+^, Biowest). The BRET^2^-ratio was normalized by defining the mean values for the RLuc8 signal as 0% and the signal prior to addition of the cAMP stimuli as 100%.

### Expression, purification of recombinant Cα and His-Riβ

The human RIβ wt and L50R constructs were expressed in *E. coli* TP2000 cells (50, 51) to obtain cAMP free R subunits. To obtain high amounts of recombinant proteins, liters of cell cultures were inoculated with colonies from transfection and incubated at 37 °C under shaking at 180 rpm till an optical density of ∼0.6 was reached. Protein expression was induced by adding 400 μM IPTG following incubation overnight (at room temperature). Purification of the wt and L50R mutant was performed using a Ni^2+^-NTA-Agarose (Protino Ni^2+^-NTA Agarose, Macherey-Nagel^TM^, DE) via their N-terminal polyhistidine Tags (His_7_-Tag). Harvested cell pellets were treated with lysis buffer containing 50 mM KH_2_PO_4_ (pH 8.0), 500 mM NaCl, 20 mM imidazole, 5 mM β-Mercaptoethanol as well as freshly added 0.1 % Triton X-100 and protease inhibitor (cOmplete ™ EDTA-free protease inhibitor cocktail, Sigma-Aldrich) and lysed via French Pressure Cell (Thermo IEC). After centrifugation at 45.000 xg for 30 min (at 4°C) supernatant was collected and applied to a previously equilibrated agarose resin. After 1 hour of incubation at 4°C resins were washed by centrifuging the samples for 2 min at 1000 xg (at 4°C) with two different wash buffers containing increasing amounts of imidazole. After two wash steps with the first wash buffer containing 50 mM KH_2_PO_4_ (pH 8.0), 500 mM NaCl, 60 mM imidazole, 5 mM β-Mercaptoethanol a last wash step was performed with the following buffer 50 mM KH_2_PO_4_ (pH 8.0), 500 mM NaCl, 100 mM imidazole, 5 mM β-Mercaptoethanol. The elution of the R subunits was performed with 50 mM KH_2_PO_4_ (pH 8.0), 500 mM NaCl, 250 mM imidazole, 5 mM β-Mercaptoethanol. Each purification step was analyzed by an SDS-Gel. Proteins were stored on ice at 4°C in 20 mM MOPS (pH 7.0), 150 mM NaCl, 2 mM EGTA, 2 mM EDTA and 5 mM β-Mercaptoethanol. Overexpression of the C subunit was performed in *E.coli* BL21(DE3) cells and purified using with IP20 affinity chromatography as already described (52, 53).

### Holoenzyme activation

Holoenzyme was preformed by mixing C- and R-subunits in a 1:1.2 ratio in 200µl for 1 h on ice at 4°C in 20 mM MOPS, pH 7.4, 100 mM NaCl, 1mM ATP, 10 mM MgCl_2_, 2 mM β-Mercaptoethanol and 0.5 mg/ml BSA. Holoenzyme activation was determined by a coupled spectrophotometric kinase assay as described by (54), where the activity of the C-subunit was measured by the phosphorylation of the synthetic peptide Kemptide (Leu-Arg-Arg-Ala-Ser-Leu-Gly) (GeneCust). Measurements were performed with the SPECORD 200 spectrophotometer (Analytic Jena). 20 nM of C-subunit and 24 nM R-subunit (final concentrations) were used in each measurement in 100 µl Cook Assay Mix (100 mM MOPS, pH 7.0, 10 mM MgCl_2_, 1 mM Phosphoenolpyruvate, 1 mM ATP, 15 U/ml Lactate dehydrogenase, 8.4 U/ml Pyruvate kinase, 5 mM β-Mercaptoethanol and 0.2 mM NADH). Different cAMP concentrations (in 1 µl) were used to induce holoenzyme dissociation and the reaction was started by adding 1 µl of Kemptide (final concentration of 251 µM). All dilutions were prepared in 20 mM MOPS, pH 7.4, 150 mM MgCl_2_, 2 mM β-Mercaptoethanol and 0.5 mg/ml BSA. To obtain the kinase activity the extinction of NADH was measured at 340 nm for 30 seconds and plotted over the time. The obtained slope values (Δ340nm/min) were then plotted against increasing cAMP concentrations. For calculation of the K_act_ values a sigmoidal dose-response fit was applied.

### Surface plasmon resonance analysis

Interaction of RIβ-WT or RIβ-L50R with an FSS -tagged hCα subunit was analyzed by Surface plasmon resonance (SPR) using a Biacore T200 (Cytiva life sciences, UK). FSS-hCα was captured as ligand on a Biacore CM5-Chip T200 (Cytiva life sciences, UK) previously coated with Strep-Tactin®XT using the Twin-Strep-tag® Capture Kit of IBA Lifesciences GmbH (Göttingen, DE) as described by the manual. N-terminal His tagged RIβ-subunits were used as analytes. All SPR measurements were performed in 10 mM Hepes, 150 mM NaCl, 1 mM ATP, 10 mM MgCl_2_ and 0.005 % Tween20. One flow cells of the chip was used as reference. On the reference cell no catalytic subunit was immobilized. Capturing of 5 nM C subunit on the measurement cell (second flow cell) was performed for 50 s at a flow rate of 10 μl/min. After capturing a series of R concentrations were applied at a flow rate of 30 µl/min to both flow cells. After 300 s association, injection of buffer without R subunit (30 µl/min flow rate) was performed to start dissociation (300s). As multi-cycle kinetics were performed, regeneration of the chip surface for each R concentration was induced by treating the chip three times with a 3 M Guanidinium chloride (GuHCl) solution for 60 s at a flow rate of 30 μl/min. For all the SPR measurements functional concentration of R subunit was used. For the determination of the functional concentration a fixed concentration of 5 nM C subunit was titrated with different concentrations of cAMP-free R subunits. As indicator for C-subunit activity, phosphorylation of the synthetic peptide Kemptide was measured with the spectrophotometric assay an in (54). For analyzing the binding kinetics of C- and R-subunit the plotted SPR-curves were fitted with a 1:1 Langmuir binding model (A + B ↔ AB) for calculation of association (k_ass_) and dissociation (k_diss_) rates as well as for the equilibrium dissociation constant (K_D_). All measurements were performed on a Biacore® T200 device (Cytiva life sciences, UK). Data evaluation was performed with the software Biacore T200 evaluation software 3.2 (Cytiva life sciences, UK).

To test the cAMP-triggered dissociation of the RIβ-WT or RIβ-L50R from the catalytic subunit FSS-hCa was captured to a Strep-Tactin®XT coated CM5-Chip as described above. Wt or mutant R subunit (15 nM) were injected over the immobilized FSS-C subunit for 300 s and holoenzyme stability was monitored for 200 s in running buffer. Subsequently increasing concentrations of cAMP were injected in the dissociation phase for 150 s to induce holoenzyme dissociation. After each cycle (capture of FSS C binding of 15nM R-subunit holoenzyme stability check cAMP triggered holoenzyme dissociation), chip regeneration was performed as described above. Data were normalized before R-subunit association (0%) and before cAMP-triggered dissociation (100%). The evaluation was performed with the software BIAevaluation 4.1.1., Biacore T200 evaluation software 3.2 (Cytiva life sciences) and GraphPad Prism 8.0.1 (GraphPad Software, Inc).

### Size-exclusion chromatography

To investigate dimer formation size-exclusion chromatography (SEC) was performed with recombinant N-terminal His-tagged RIβ-WT or RIβ-L50R using an ÄKTA go protein purification system with a Superose™ 6 Increase 3.2/300 (Cytiva life sciences, UK). Proteins were injected on the column using 25 µl or 50 µl injection loops at a flow rate of 0.04 ml/min in 20 mM MOPS (pH 7.4), 150 mM NaCl, and 2 mM β-Mercaptoethanol. The calculation of the Stokeś radii was performed using a calibration curve. Protein standards from the Gel Filtration Calibration Kit LMW of Cytiva life sciences (UK) were used. Fractions of the protein peaks of the L50R were collected and identified by Western-Blot. Detection was achieved by detecting the fluorescent signal at 800 nm using the LI-COR odyssey® Fc device (LI-COR Biosciences).

### Fluorescence polarization

The fluorescence polarization competition assay was performed by mixing in a 1:1 ratio (total volume of 60 µl) increasing cAMP concentrations, diluted in a buffer containing 8- (2-[fluoresceinyl]aminoethylthio)-cAMP (8-Fluo-cAMP) (final concentration 0.5 nM) with WT or L50R mutant protein (final concentration 4 nM) and measuring the amount of polarized light, as previously described (55). For measurements a 384-Well BRAND plates (BRAND) was used and each cAMP concentration was measured in duplicates. For the detection of the intensity of polarized light, a CLARIOstar Omega plate reader (BMG LABTECH) was used. All dilutions were performed in a buffer containing 20 mM MOPS, pH 7.4, 150 mM NaCl, 0.005% CHAPS and 1 mM DTT. Data were blank subtracted using the MARS evaluation software (BMG LABTECH) and analyzed to obtain EC_50_ values.

## Supplementary figures

**Figure. S1.**
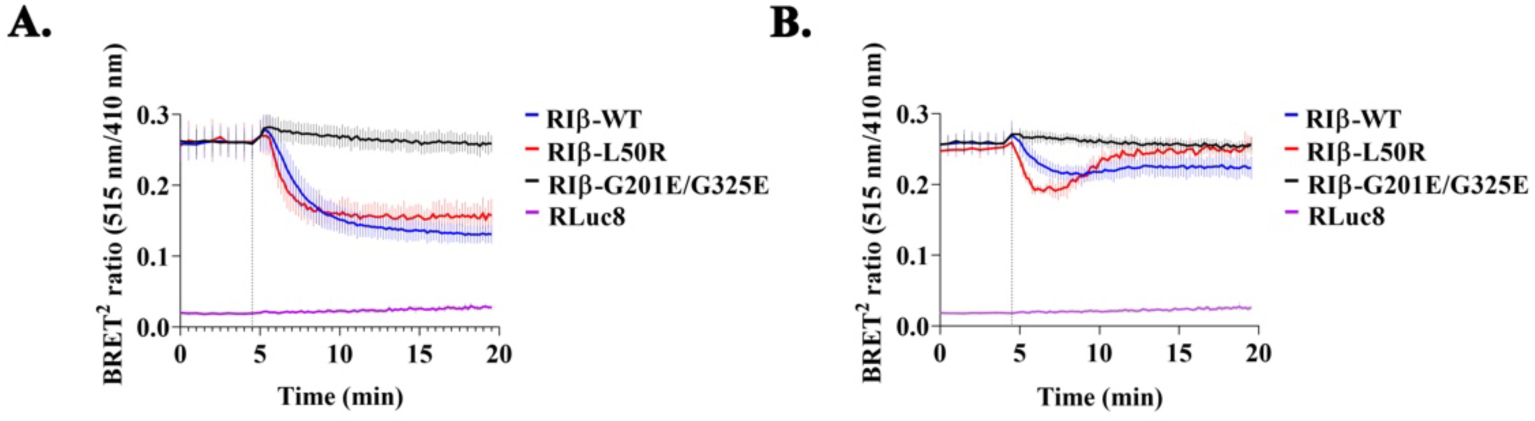
Raw data for time-dependent BRET^2^ measurements. HEK293 cells were stimulated with 50 µM FSK and 100 µM IBMX (A) or with 1 µM ISO (B). Injection times of the respective stimuli are indicated with a dotted line. Data represent the BRET^2^ ratio corresponding to the GFP^2^-hCα signal (515 nm) over a hRIβ-RLuc8 signal (410 nm) for RIβ-WT and RIβ-L50R. The signal for the RIβ double G201E/G325E unable to bind cAMP was used as a control for no RIβ:C dissociation. The signal from protein encoding by a Rluc8 empty vector was used for data normalization, as presented in Fig. 4. Each curve represents the mean of six replicates ± SEM.

**Figure. S2.**
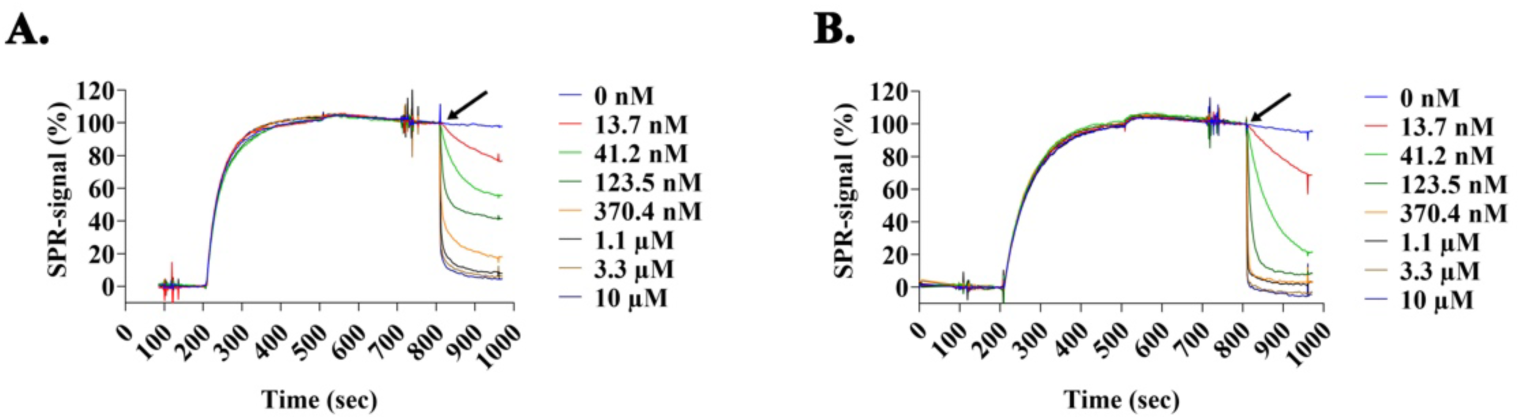
SPR analysis was used to determine cAMP-induced dissociation of RIβ-WT:Cα or RIβ-L50R:Cα. SRP sensograms of FSS-Cα subunit bound to a Strep-Tactin-coated CM5 sensor chip. Fifteen nM RIβ-WT (A) or RIβ-L50R (B) were injected and associated to the FSS-Cα subunit, as indicated by the increased SRP signal. Different concentrations of cAMP (black arrows) induced an increasing rate of RIβ:C dissociation, as indicated by reduction of the SRP signal. The data were normalized at approximately 190 sec (0%) and 800 sec (100%). Zoomed in sensograms showing the cAMP-induced dissociation of the RIβ-WT:Cα or RIβ-L50R:Cα holoenzymes are presented in Figure 6 D-E.

## References

1. Cadd, G. and McKnight, G.S. (1989) Distinct patterns of cAMP-dependent protein kinase gene expression in mouse brain. Neuron, 3, 71–79.

2. Kandel, E.R. (2012) The molecular biology of memory: cAMP, PKA, CRE, CREB-1, CREB-2, and CPEB. Mol Brain, 5, 14.

3. Abel, T., Nguyen, P. V, Barad, M., Deuel, T.A., Kandel, E.R. and Bourtchouladze, R. (1997) Genetic demonstration of a role for PKA in the late phase of LTP and in hippocampus-based long-term memory. Cell, 88, 615–626.

4. Huang, Y.Y., Kandel, E.R., Varshavsky, L., Brandon, E.P., Qi, M., Idzerda, R.L., McKnight, G.S. and Bourtchouladze, R. (1995) A genetic test of the effects of mutations in PKA on mossy fiber LTP and its relation to spatial and contextual learning. Cell, 83, 1211–1222.

5. Ma, L., Day-Cooney, J., Benavides, O.J., Muniak, M.A., Qin, M., Ding, J.B., Mao, T. and Zhong, H. (2022) Locomotion activates PKA through dopamine and adenosine in striatal neurons. Nature 2022 611:7937, 611, 762–768.

6. Dagda, R.K. and Banerjee, T. Das (2015) Role of PKA in regulating mitochondrial function and neuronal development: implications to neurodegenerative diseases. Rev Neurosci, 26, 359.

7. Da Cruz, A.B., Wentzell, J. and Kretzschmar, D. (2008) Swiss Cheese, a Protein Involved in Progressive Neurodegeneration, Acts as a Noncanonical Regulatory Subunit for PKA-C3. Journal of Neuroscience, 28, 10885–10892.

8. Licata, N. V, Cristofani, R., Salomonsson, S., Wilson, K.M., Kempthorne, L., Vaizoglu, D., D’Agostino, V.G., Pollini, D., Loffredo, R., Pancher, M., et al. (2022) C9orf72 ALS/FTD dipeptide repeat protein levels are reduced by small molecules that inhibit PKA or enhance protein degradation. EMBO J, 41.

9. Jicha, G.A., Weaver, C., Lane, E., Vianna, C., Kress, Y., Rockwood, J. and Davies, P. (1999) cAMP-Dependent Protein Kinase Phosphorylations on Tau in Alzheimer’s Disease. Journal of Neuroscience, 19, 7486–7494.

10. Carlyle, B.C., Nairn, A.C., Wang, M., Yang, Y., Jin, L.E., Simen, A.A., Ramos, B.P., Bordner, K.A., Craft, G.E., Davies, P., et al. (2014) CAMP-PKA phosphorylation of tau confers risk for degeneration in aging association cortex. Proc Natl Acad Sci U S A, 111, 5036–5041.

11. Greggio, E., Bubacco, L. and Russo, I. (2017) Cross-talk between LRRK2 and PKA: implication for Parkinson’s disease? Biochem Soc Trans, 45, 261–267.

12. Muda, K., Bertinetti, D., Gesellchen, F., Hermann, J.S., Von Zweydorf, F., Geerlof, A., Jacob, A., Ueffing, M., Gloeckner, C.J. and Herberg, F.W. (2014) Parkinson-related LRRK2 mutation R1441C/G/H impairs PKA phosphorylation of LRRK2 and disrupts its interaction with 14-3-3. Proc Natl Acad Sci U S A, 111.

13. Palomo, V., Nozal, V., Rojas-Prats, E., Gil, C. and Martinez, A. (2021) Protein kinase inhibitors for amyotrophic lateral sclerosis therapy. Br J Pharmacol, 178, 1316–1335.

14. Lin, J.-T., Chang, W.-C., Chen, H.-M., Lai, H.-L., Chen, C.-Y., Tao, M.-H. and Chern, Y. (2013) Regulation of Feedback between Protein Kinase A and the Proteasome System Worsens Huntington’s Disease. Mol Cell Biol, 33, 1073.

15. Omar, M.H. and Scott, J.D. (2020) AKAP Signaling Islands: Venues for Precision Pharmacology. Trends Pharmacol Sci, 41, 933–946.

16. Taylor, S.S., Ilouz, R., Zhang, P. and Kornev, A.P. (2012) Assembly of allosteric macromolecular switches: lessons from PKA. Nat Rev Mol Cell Biol, 13, 646–658.

17. Ilouz, R., Lev-Ram, V., Bushong, E.A., Stiles, T.L., Friedmann-Morvinski, D., Douglas, C., Goldberg, G., Ellisman, M.H. and Taylor, S.S. (2017) Isoform-specific subcellular localization and function of protein kinase A identified by mosaic imaging of mouse brain. Elife, 6.

18. Zhang, P., Smith-Nguyen, E. V, Keshwani, M.M., Deal, M.S., Kornev, A.P. and Taylor, S.S. (2012) Structure and allostery of the PKA RIIbeta tetrameric holoenzyme. Science (1979), 335, 712–716.

19. Marbach, F., Stoyanov, G., Erger, F., Stratakis, C.A., Settas, N., London, E., Rosenfeld, J.A., Torti, E., Haldeman-Englert, C., Sklirou, E., et al. (2021) Variants in PRKAR1B cause a neurodevelopmental disorder with autism spectrum disorder, apraxia, and insensitivity to pain. Genet Med, 23, 1465–1473.

20. Wong, T.H., Chiu, W.Z., Breedveld, G.J., Li, K.W., Verkerk, A.J., Hondius, D., Hukema, R.K., Seelaar, H., Frick, P., Severijnen, L.A., et al. (2014) PRKAR1B mutation associated with a new neurodegenerative disorder with unique pathology. Brain, 137, 1361–1373.

21. Sanderson, J.L., Freund, R.K., Gorski, J.A. and Dell’Acqua, M.L. (2021) β-Amyloid disruption of LTP/LTD balance is mediated by AKAP150-anchored PKA and Calcineurin regulation of Ca2+-permeable AMPA receptors. Cell Rep, 37, 109786.

22. Sarma, G.N., Kinderman, F.S., Kim, C., von Daake, S., Chen, L., Wang, B.C. and Taylor, S.S. (2010) Structure of D-AKAP2:PKA RI complex: insights into AKAP specificity and selectivity. Structure, 18, 155–166.

23. Dahlin, H.R., Zheng, N. and Scott, J.D. (2021) Beyond PKA: Evolutionary and structural insights that define a docking and dimerization domain superfamily. Journal of Biological Chemistry, 297, 100927.

24. Drougat, L., Settas, N., Ronchi, C.L., Bathon, K., Calebiro, D., Maria, A.G., Haydar, S., Voutetakis, A., London, E., Faucz, F.R., et al. (2021) Genomic and sequence variants of protein kinase A regulatory subunit type 1β (PRKAR1B) in patients with adrenocortical disease and Cushing syndrome. Genetics in Medicine, 23, 174–182.

25. Benjamin-Zukerman, T., Shimon, G., Gaine, M.E., Dakwar, A., Peled, N., Aboraya, M., Masri-Ismail, A., Safadi-Safa, R., Solomon, M., Lev-Ram, V., et al. (2024) A mutation in the PRKAR1B gene drives pathological mechanisms of neurodegeneration across species. Brain, 10.1093/BRAIN/AWAE154.

26. Kim, C., Cheng, C.Y., Saldanha, S.A. and Taylor, S.S. (2007) PKA-I Holoenzyme Structure Reveals a Mechanism for cAMP-Dependent Activation. Cell, 130, 1032–1043.

27. Woodford, T.A., Correll, L.A., McKnight, G.S. and Corbin, J.D. (1989) Expression and Characterization of Mutant Forms of the Type I Regulatory Subunit of cAMP-dependent Protein Kinase: The effect of defective Camp binding on holoenzyme activation. Journal of Biological Chemistry, 264, 13321–13328.

28. Steinberg, R.A., O’Farrell, P.H., Friedrich, U. and Coffino, P. (1977) Mutations causing charge alterations in regulatory subunits of the cAMP-dependent protein kinase of cultured S49 lymphoma cells. Cell, 10, 381–391.

29. Angers, S., Salahpour, A., Joly, E., Hilairet, S., Chelsky, D., Dennis, M. and Bouvier, M. (2000) Detection of β2-adrenergic receptor dimerization in living cells using bioluminescence resonance energy transfer (BRET). Proc Natl Acad Sci U S A, 97, 3684–3689.

30. Lands, A.M., Luduena, F.P. and Buzzo, H.J. (1967) Differentiation of receptors responsive to isoproterenol. Life Sci, 6, 2241–2249.

31. Taylor, S.S., Wallbott, M., MacHal, E.M.F., Søberg, K., Ahmed, F., Bruystens, J., Vu, L., Baker, B., Wu, J., Raimondi, F., et al. (2021) PKA Cβ: a forgotten catalytic subunit of cAMP-dependent protein kinase opens new windows for PKA signaling and disease pathologies. Biochemical Journal, 478, 2101–2119.

32. Herberg, F.W., Taylor, S.S., Dostmann, W.R.G., Zorn, M. and Davis, S.J. (1994) Crosstalk between Domains in the Regulatory Subunit of cAMP-Dependent Protein Kinase: Influence of Amino Terminus on cAMP Binding and Holoenzyme Formation. Biochemistry, 33, 7485–7494.

33. Bonifati, V., Rizzu, P., Van Baren, M.J., Schaap, O., Breedveld, G.J., Krieger, E., Dekker, M.C.J., Squitieri, F., Ibanez, P., Joosse, M., et al. (2003) Mutations in the DJ-1 gene associated with autosomal recessive early-onset parkinsonism. Science (1979), 299, 256–259.

34. Baulac, S., LaVoie, M.J., Strahle, J., Schlossmacher, M.G. and Xia, W. (2004) Dimerization of Parkinson’s disease-causing DJ-1 and formation of high molecular weight complexes in human brain. Molecular and Cellular Neuroscience, 27, 236–246.

35. Moore, D.J., Zhang, L., Dawson, T.M. and Dawson, V.L. (2003) A missense mutation (L166P) in DJ-1, linked to familial Parkinson’s disease, confers reduced protein stability and impairs homo-oligomerization. J Neurochem, 87, 1558–1567.

36. Berdyński, M., Miszta, P., Safranow, K., Andersen, P.M., Morita, M., Filipek, S., Żekanowski, C. and Kuźma-Kozakiewicz, M. (2022) SOD1 mutations associated with amyotrophic lateral sclerosis analysis of variant severity. Sci Rep, 12, 103.

37. Wang, J., Caruano-Yzermans, A., Rodriguez, A., Scheurmann, J.P., Slunt, H.H., Cao, X., Gitlin, J., Hart, P.J. and Borchelt, D.R. (2007) Disease-associated mutations at copper ligand histidine residues of superoxide dismutase 1 diminish the binding of copper and compromise dimer stability. Journal of Biological Chemistry, 282, 345–352.

38. Basith, S., Manavalan, B. and Lee, G. (2022) Amyotrophic lateral sclerosis disease-related mutations disrupt the dimerization of superoxide dismutase 1 - A comparative molecular dynamics simulation study. Comput Biol Med, 151, 106319.

39. Ilouz, R., Bubis, J., Wu, J., Yim, Y.Y., Deal, M.S., Kornev, A.P., Ma, Y., Blumenthal, D.K. and Taylor, S.S. (2012) Localization and quaternary structure of the PKA RIbeta holoenzyme. Proc Natl Acad Sci U S A, 109, 12443–12448.

40. Zhang, B., Gaiteri, C., Bodea, L.G., Wang, Z., McElwee, J., Podtelezhnikov, A.A., Zhang, C., Xie, T., Tran, L., Dobrin, R., et al. (2013) Integrated systems approach identifies genetic nodes and networks in late-onset Alzheimer’s disease. Cell, 153, 707.

41. James Surmeier, D., Obeso, J.A. and Halliday, G.M. (2017) Parkinson’s Disease Is Not Simply a Prion Disorder. Journal of Neuroscience, 37, 9799–9807.

42. Puigdellívol, M., Saavedra, A. and Pérez-Navarro, E. (2016) Cognitive dysfunction in Huntington’s disease: mechanisms and therapeutic strategies beyond BDNF. Brain Pathology, 26, 752–771.

43. Mead, R.J., Shan, N., Reiser, H.J., Marshall, F. and Shaw, P.J. (2023) Amyotrophic lateral sclerosis: a neurodegenerative disorder poised for successful therapeutic translation. Nat Rev Drug Discov, 22, 185–212.

44. De, A., Loening, A.M. and Gambhir, S.S. (2007) An Improved Bioluminescence Resonance Energy Transfer Strategy for Imaging Intracellular Events in Single Cells and Living Subjects. Cancer Res, 67, 7175–7183.

45. Micsonai, A., Moussong, É., Wien, F., Boros, E., Vadászi, H., Murvai, N., Lee, Y.H., Molnár, T., Réfrégiers, M., Goto, Y., et al. (2022) BeStSel: webserver for secondary structure and fold prediction for protein CD spectroscopy. Nucleic Acids Res, 50, W90–W98.

46. Micsonai, A., Moussong, É., Murvai, N., Tantos, Á., Tőke, O., Réfrégiers, M., Wien, F. and Kardos, J. (2022) Disordered–Ordered Protein Binary Classification by Circular Dichroism Spectroscopy. Front Mol Biosci, 9, 863141.

47. Micsonai, A., Bulyáki, É. and Kardos, J. (2021) BeStSel: From Secondary Structure Analysis to Protein Fold Prediction by Circular Dichroism Spectroscopy. Methods in Molecular Biology, 2199, 175–189.

48. Micsonai, A., Wien, F., Bulyáki, É., Kun, J., Moussong, É., Lee, Y.H., Goto, Y., Réfrégiers, M. and Kardos, J. (2018) BeStSel: a web server for accurate protein secondary structure prediction and fold recognition from the circular dichroism spectra. Nucleic Acids Res, 46, W315– W322.

49. Micsonai, A., Wien, F., Kernya, L., Lee, Y.H., Goto, Y., Réfrégiers, M. and Kardos, J. (2015) Accurate secondary structure prediction and fold recognition for circular dichroism spectroscopy. Proc Natl Acad Sci U S A, 112, E3095–E3103.

50. Roy, A. and Danchin, A. (1981) Restriction map of the cya region of the Escherichia coli K12 chromosome. Biochimie, 63, 719–722.

51. Kim, J.J., Flueck, C., Franz, E., Sanabria-Figueroa, E., Thompson, E., Lorenz, R., Bertinetti, D., Baker, D.A., Herberg, F.W. and Kim, C. (2015) Crystal Structures of the Carboxyl cGMP Binding Domain of the Plasmodium falciparum cGMP-dependent Protein Kinase Reveal a Novel Capping Triad Crucial for Merozoite Egress. PLoS Pathog, 11, e1004639.

52. Zhang, P., Knape, M.J., Ahuja, L.G., Keshwani, M.M., King, C.C., Sastri, M., Herberg, F.W. and Taylor, S.S. (2015) Single Turnover Autophosphorylation Cycle of the PKA RIIβ Holoenzyme. PLoS Biol, 13, e1002192.

53. Affinity purification of the C alpha and C beta isoforms of the catalytic subunit of cAMP-dependent protein kinase - PubMed.

54. Cook, P.F., Neville Jr., M.E., Vrana, K.E., Hartl, F.T. and Roskoski Jr., R. (1982) Adenosine cyclic 3’, 5’-monophosphate dependent protein kinase: kinetic mechanism for the bovine skeletal muscle catalytic subunit. Biochemistry, 21, 5794–5799.

55. Moll, D., Prinz, A., Gesellchen, F., Drewianka, S., Zimmermann, B. and Herberg, F.W. Biomolecular interaction analysis in functional proteomics. 10.1007/s00702-006-0515-5.

